# Localised activity of reverse gyrase at gene regulatory elements

**DOI:** 10.64898/2025.12.08.692955

**Authors:** Paul Villain, Vladislav Kuzin, Augustin Darennes-Degaugue, Florence Lorieux, Antoine Hocher, Romain Le Bars, Tobias Warnecke, Laura Baranello, Tamara Basta

## Abstract

DNA topoisomerases are essential enzymes found in all cells, where they regulate DNA supercoiling. Reverse gyrase (RG) is a unique type of topoisomerase that introduces positive supercoils into DNA and appears exclusively in hyperthermophiles where it was proposed to play a key, yet still elusive, role. Here, we investigate RG activity in the hyperthermophilic archaeon *Thermococcus kodakarensis* at 85°C, its optimal growth temperature, using genetics and functional genomics assays. Deletion of RG led to a loss of positive supercoiling in plasmid DNA and the reduced dynamic range of transcription, without affecting histone occupancy. To investigate the effects of RG loss on the topology of chromosomal DNA, we established a psoralen photobinding assay (TMP-seq) in *T. kodakarensis* under native growth conditions. TMP enrichment patterns were consistent with the twin-domain model of transcription and further revealed that promoters of expressed transcription units are, on average, underwound. TMP-seq profiles in an RG deletion strain revealed that promoters are hotspots for RG activity, consistent with RG acting on hyper-negatively supercoiled substrates. We propose that RG acts not as a global modulator of supercoiling, but as a local genome guardian, selectively stabilising vulnerable regulatory regions to ensure a delicate balance between DNA accessibility and integrity under extremely high temperatures.

## INTRODUCTION

DNA topology is a direct consequence of the double-helical nature of DNA, and is defined by how the two complementary DNA strands are intertwined. The most commonly found topology in cells is DNA supercoiling whereby the DNA double helix winds around itself to form a superhelix that can be positively (left-handed) or negatively (right-handed) overwound. Negatively supercoiled DNA is easier to denature while positively supercoiled DNA is more stable and compact^1^. Consequently, these two conformational states differ in how they affect DNA transactions that require the opening of the double helix (e.g. replication, transcription, repair) as well as higher order chromosome folding^1,2^, therefore affecting pivotal biological processes.

The deleterious effects of high temperature on DNA are well documented, with a roughly 3,000-fold increase in DNA degradation for growth at 100°C compared to 37°C^3^. The question of how hyperthermophiles, microorganisms that thrive at temperatures above 80°C, maintain the integrity of their DNA at such extreme temperatures has captivated the scientific community for decades. Hyperthermophilic archaea are the only organisms that naturally harbour positively supercoiled DNA as shown by early studies of plasmids isolated from these organisms^4^. Positive supercoiling activity was first identified in 1984 in cellular extracts of the acidothermophilic archaeon *Sulfolobus acidocaldarius* and was linked to the presence of a unique topoisomerase, reverse gyrase (RG)^5^. RG is an atypical topoisomerase formed by the fusion of a RecQ-like helicase domain to a *bona fide* Type IA topoisomerase^6^. *In vitro* characterisation of this enzyme showed that it catalysed positive supercoiling either from strongly negatively supercoiled DNA, from relaxed plasmid DNA or from DNA containing single-stranded bubbles in an ATP-dependent fashion whereby the coupling of the helicase and topoisomerase activities are required to generate one positive supercoil per catalytic cycle^7–10^.

RG is found systematically in hyperthermophiles and most thermophiles but never in mesophiles, suggesting that it has a specific and critical role for sustaining life at high temperature^11,12^. Surprisingly, the RG-encoding gene, *rgy*, can be deleted in the hyperthermophilic archaea *Thermococcus kodakarensis* and *Pyrococcus furiosus*. However, deletion mutants could not grow at temperatures above 95°C, suggesting that RG is indeed essential for viability when cells are exposed to extreme thermal conditions^13,14^. In the acidothermophile *Saccharolobus islandicus*, which grows optimally at 75°C, the *topR2* gene, which encodes one of the two RG paralogs present in this organism, could not be deleted indicating that in this organism RG is critically important at much lower temperature^15,16^.

Several studies reported the involvement of RG in maintaining genome integrity and in DNA repair^17^. RG was shown to protect DNA from breaks *in vivo*^18^, to be recruited to UV-damaged DNA^19^, to protect DNA from degradation^10,20^ independently of its topoisomerase activity, and to inhibit the activity of translesion DNA polymerase *in vitro*^21^. Other studies reported the increase of positive supercoiling of plasmids following heat shock or UV irradiation suggesting that the positive supercoiling activity of the RG may be activated under conditions favouring DNA damage^4,22,23^.

While significant progress has been made in the last four decades to understand the *in vitro* properties of RG, the functional significance of its positive supercoiling activity *in vivo* remains unclear^24^. To address this issue, genome-wide mapping of supercoiled DNA is needed. Trimethylpsoralen (TMP) photobinding has been used since the 1980s to map supercoiling in bacterial and eukaryotic model organisms^25–27^, and more recently, in combination with next generation sequencing, to map supercoiling with nucleotide-scale precision^28–30^. Psoralen is a small planar aromatic molecule that preferentially intercalates into underwound (negatively supercoiled) DNA and can establish covalent interstrand crosslinks under UV light^25^. Crosslinked DNA can then be separated from non-crosslinked DNA on a denaturing agarose gel and sequenced. The Crosslinking Level (CLL), defined as the log2 ratio of crosslinked to non-crosslinked DNA, then provides a quantitative picture of psoralen intercalation across the genome^27,31^.

In this work, we used a genetic approach in combination with genome-wide assays, including TMP photobinding, to investigate the contribution of RG to DNA supercoiling and gene expression *in vivo* on a genome-wide scale. RG deletion led to widespread deregulation of transcription characterized by the compression of the dynamic range of gene expression. We explain this phenotype by demonstrating that removal of RG affects supercoiling levels at promoters, while leaving histone occupancy unaffected. Our findings challenge the prevailing model that the primary function of RG is to introduce pervasive positive supercoiling into the chromosomes of hyperthermophiles. Instead, they reinforce the idea that RG specifically targets underwound, AT-rich, and histone-depleted DNA—such as promoters—which are particularly prone to destabilization and damage at elevated temperatures.

## MATERIALS AND METHODS

### Construction and culturing of recombinant *Thermococcus kodakarensis* strains

Strains, plasmids and oligonucleotides used in this study are listed in **Table S1**. Plasmids for genetic modification of *T. kodakarensis* were constructed using standard cloning procedures and amplified in *E. coli* XL1-Blue (Agilent) grown on LB medium supplemented with Ampicillin (100 µg/ml), Kanamycin (50 µg/ml) or Chloramphenicol (33 µg/ml).

*Thermococcus* strains used in this study are derivatives of *T. kodakarensis* KOD1 (RefSeq NC_006624.1) and were grown heterotrophically in strict anaerobic conditions in Ravot medium^32^ containing sulphur as the final electron acceptor. To monitor growth, liquid cultures were inoculated with overnight precultures at 1:100 dilution and cell density was determined using a Thoma cell counting chamber (0.01 mm depth) and phase-contrast microscopy. The generation time (g) was calculated using the equation g = ln(2)/µ where µ is the specific growth rate in min^-1^, defined as the maximum slope of the ln transformation of the growth curve.

Transformation of *T. kodakarensis* cells was performed as previously described^33^. The *T. kodakarensis* DAD and DADΔRG strains (kindly provided by Hiroki Higashibata, Toyo University, **Table S1**), are both auxotroph for uracil (Δ*pyrF*) and agmatine (Δ*TK0149*). DADΔRG carries, in addition, a markerless deletion of the gene encoding reverse gyrase (Δ*TK0470*). A subsequent Δ*trpE*::*pyrF* replacement was made in these strains to allow tryptophan-based selection, resulting in TKDAD and TKΔRG strains. These two strains were transformed with a small replicating plasmid, pTPTK3, carrying the *pdaD* gene^34^ resulting in the TKDADp and TKΔRGp strains. For simplicity, henceforth, TKDADp and TKΔRGp strains will be denominated WT and ΔRG strains, respectively.

### Confocal microscopy and image analysis

Exponentially growing or stationary phase cultures of *T. kodakarensis* were fixed with 1.2 % (v/v) final concentration of formaldehyde for 15 min at 85°C with shaking. Formaldehyde crosslinking was quenched with 250 mM of glycine and 500 µL of culture was centrifuged (5000 g) for 5 min at room temperature. The cell pellet was resuspended in 1 ml of 1× Ravot salt solution containing 1 µg/ml of DAPI and incubated in the dark for 15 min. The cells were then centrifuged (5000 g) for 5 min at room temperature, the cell pellet was washed once with 250 µL of 1× Ravot salt solution and resuspended in 25 µL of 1× Ravot salt solution. Cell suspensions (2 µL) were mounted on glass slides covered with a thin layer of 1% (w/v) agarose dissolved in water. Differential interference contrast (DIC) and fluorescent images were obtained at room temperature on an SP8 confocal laser scanning microscope (Leica Microsystems) equipped with hybrid detectors and a 63x oil immersion objective (HC Plan Apo, 1.4 numerical aperture; Leica). Fluorescence detection was performed by exciting the sample with a 405 nm laser and collecting fluorescence between 415 and 515 nm at the speed of 600 Hz with a line averaging of three. The image format was adjusted to provide an XY optimal sampling (pixel size of 60 nm) and for each position z-stacks (3 µm width and 0.5 µm step) were acquired.

To perform automated cell detection, confocal stacks were initially projected along the z-axis using a maximal intensity projection algorithm. The 2D images obtained were then converted into masks using the pre-trained model “bact_fluor_omni” of Omnipose^35^, and the objects within the masks were eroded (outermost pixel row was removed from the mask of each cell) and multiplied with the original intensity image to obtain individual objects with intensity information. Finally, MicrobeJ ^36^ with the default thresholding method was used to detect individual cells after applying morphometric filters (cell area: 0.3 - 8 µm² / length: 0.5 – 5 µm / width: 0.5 – 3 µm / cell circularity > 0.7 ) and perform cell measurements. For measurements of cell and nucleoid size, circularity, and area, at least 10 independent images were analysed per strain, totalling at least 1000 cells.

### Two-dimensional agarose gel electrophoresis

pTPTK3 DNA (4.7 kbp) was isolated from late exponential phase cultures of WT and ΔRG strains using a previously published protocol^33^. Two-dimensional agarose gel electrophoresis was performed as previously described^33^ with minor modifications. Briefly, agarose gels were prepared by dissolving 0.8 % (w/v) of agarose in 1X TBE (89 mM Tris, 89 mM boric acid, 2 mM EDTA). Electrophoresis was performed at room temperature (usually 20°C) and a minimum of 2 µg of plasmid DNA was used for each sample. In the first dimension, no intercalating agent or 1.5 µg/ml of chloroquine was added in the gel and in the running buffer (1X TBE). Electrophoresis was run at 1.2 V/cm for 22 h. Gels were subsequently equilibrated for 1 h in 1X TBE buffer supplemented with 7.5 µg/ml of chloroquine and placed in the tray after a clockwise rotation of 90°. In the second dimension, an electric field of 2 V/cm was applied for 5 h. Gels were washed 3 x 30 min in water to remove chloroquine and then stained for 1h with 2.5 µg/ml of ethidium bromide. Gels were then rinsed with water and imaged with a Typhoon Imager (Cytiva Life Sciences) using the Cy3 channel. The topoisomers were quantified from gel images by measuring the intensity of plasmid bands using the Fiji software^37^. The major topoisomer was used for the calculation of the superhelical density (σ) according to the formula σ = Wr/Tw where Wr is the number of supercoils and Tw is the number of helical turns which can be calculated by dividing the number of bp contained in the plasmid (4.7 kbp) by 10.5 which is the number of bp per helix turn in B-DNA. The calculations were corrected for the impact of the temperature shift (from 85°C, the optimal growth temperature of *T. kodakarensis*, to the temperature of electrophoresis, ∼22°C) according to the formula – 0.011°/°C/bp^38^ and for the impact of chloroquine when necessary.

### *In vivo* DNA crosslinking with psoralen under optimal growth conditions

The protocol for psoralen crosslinking of *T. kodakarensis* DNA is based on the protocol published by Kouzine and collaborators^31^. Trimethylpsoralen (TMP) (Sigma-Aldrich, ref. T6137) was diluted to saturation (0.9 mg/ml) in absolute ethanol.

For initial screening of appropriate chromosomal DNA crosslinking conditions increasing concentrations of TMP or equivalent volumes of ethanol (negative control), were injected into sealed bottles containing 20 ml of late exponential phase *T. kodakarensis* cultures (TKgyrAB strain, **Table S1**). The concentration of 1 µg/ml was chosen for preparing samples for deep sequencing. After 10 min of incubation at 85°C with TMP, cultures were poured into the lid of a 10 cm glass Petri dish maintained at 85°C using a water-bath. Cells were immediately irradiated with ∼9.6 kJ.m^-2^.min^-1^ of UV light (λ = 365 nm) using a 45 W lamp (Vilber Lourmat model VL-315.BL) for 3 min (28.8 kJ/m^2^ dose in total). The anaerobic state of cultures during irradiation was controlled through the resazurin indicator contained in the Ravot medium. After UV-irradiation, cells were immediately chilled on ice and harvested at 4°C by 15 min of centrifugation at 5,000 g.

### Total DNA extraction and fragmentation

Pellets of UV-crosslinked cells were resuspended in 250 µL of TEN (40 mM Tris-HCl pH 7.5, 1 mM EDTA, 150 mM NaCl) and then lysed by the addition of 250 µL TENST (40 mM Tris-HCl pH 7.5, 1 mM EDTA, 150 mM NaCl, 1.6% N-lauryl sarcosine, 0.12% Triton X-100). Cell lysates were treated with 0.5 mg of RNAse A (Qiagen) for 1 h at 37°C and then incubated overnight at 55°C with 0.5 mg of proteinase K (Thermofisher Scientific). DNA was extracted using the classical phenol/chloroform method, with three rounds of phenol/chloroform/isoamyl alcohol (25:24:1) extraction (Sigma-Aldrich) and one round of chloroform/isoamyl alcohol extraction (VWR, Ready-Red). The aqueous phase was recovered and DNA precipitated by addition of 0.8 volume of isopropanol. After 1 h of incubation at -80°C, DNA was pelleted at 20,000 g, 4°C for 30 min. Dried pellets were resuspended in 85 µL of 5 mM Tris/HCl, pH 8.5. DNA was fragmented to an average size of 200 bp by 200 cycles of 180 s of sonication with 10% duty and 175 peak, using a S220 focused-ultrasonicator (Covaris).

### Separation of TMP crosslinked and non-crosslinked DNA fragments

The sonicated DNA was separated on a 0.65% (w/v) agarose gel in 0.5X TAE buffer. DNA fragments within the range of 100-300 bp were gel extracted with the NucleoSpin Gel and PCR Clean-up kit (Macherey-Nagel) and eluted with 15 µL of ultrapure water adjusted to pH 8.5 with sodium hydroxide. DNA was not exposed to UV light during the gel excision. After addition of 4 µL of 0.1 M phosphate buffer (57.7 mM Na_2_HPO_4_ and 42.3 mM NaH_2_PO_4_, pH 7), DNA samples were heat-denatured for 5 min at 100°C, then 20 µL of a mix of DMSO and glyoxal (Alfa Aesar) was immediately added to a final proportion of 40% (v/v) and 4% (v/v), respectively, followed by 90 min of incubation at 55°C to denature non-crosslinked DNA. Crosslinked and non-crosslinked DNA fragments were separated by electrophoresis at 2 V/cm for 14 h, with buffer recirculation between electrodes, on 3% (w/v) agarose gels. Gels were 20 cm long and made of 1.2:1.8 mixture of UltraPure agarose and NuSieve 3:1 agarose in 200 ml of 10 mM phosphate buffer pH 7. TMP-DNA crosslinks were reversed by incubating gels 3 h at 65°C in denaturing solution (0.5M NaOH, 1.5M NaCl) followed by 2 h of incubation with agitation in neutralizing solution (1.5 M NaCl, 500 mM Tris–HCl pH 8, 1 mM EDTA). Then they were rinsed in MilliQ water, equilibrated 3 times for 1 h in 1X TAE and stained overnight with SYBR Gold (Invitrogen) diluted in 1X TAE. Gels were imaged with a Typhoon Imager (Cytiva) using the Cy2 channel. Pieces of gels containing crosslinked and non-crosslinked DNA fragments were excised with a scalpel and gel-extracted using a Gel and PCR Clean-up Midi kit with NTC buffer (Macherey-Nagel). DNA was eluted with 5 mM Tris/HCl and stored at -80°C before sequencing. For each strain, biological replicates were prepared from three independent cultures. DNA-seq libraries for Illumina sequencing were constructed using the DNA SMART ChIP-Seq Kit (Takara) following the manufacturer’s protocol. Libraries were pooled in equimolar proportions and sequenced on an Illumina NextSeq500 instrument using NextSeq 500/550 High Output Kit v2 (150 cycles) and a Paired-End 2 × 75 bp run. Demultiplexing was performed with bcl2fastq2 v2.18.12. Adapters were trimmed with Cutadapt v1.15, and only reads longer than 10 bp (between 20 and 30 million reads per sample) were kept for further analysis.

### Chromatin structure profiling by micrococcal nuclease digestion

Chromatin isolation and digestion with micrococcal nuclease (MNase) were carried out as previously described^39^ with minor modifications. Briefly, recombinant *T. kodakarensis* strains were cultivated from overnight precultures in 500 ml of Ravot medium until late exponential phase. Cultures were rapidly chilled in a pre-cooled beaker immersed in a water-ice bath. Cells were pelleted by centrifugation (4500 x g, 15 min, 4°C) and washed with Ravot salt solution^32^ before storing at -80°C. Frozen pellets were resuspended in 1 ml of MNase buffer (50 mM Tris-HCl pH 8, 100 mM NaCl, and 1 mM CaCl_2_) and then subjected to 5 cycles of nitrogen freezing plus grinding with a mortar. The resulting cell lysate was recovered by pipetting and clarified by centrifugation for 5 min at 1,700 x g and 4°C. 500 µL of clarified lysates were collected by pipetting and digested for 1 h at 37°C with 70 U of RNase A (Qiagen). Samples were separated into 5 x 100 µL aliquots and then each was digested for 3 min at 37°C with 0, 50, 100, 200 or 400 U of MNase (Thermo Scientific, ≥100 U/µL). DNA from digested fractions was extracted with 300 µL of 10 mM Tris-HCl pH 8 and 400 µL of phenol/chloroform/isoamyl alcohol (25:24:1). Samples were vortexed for about 2 min and then centrifuged for 4 min at 14,000 g, room temperature. 200 µL of the top aqueous phase was collected and DNA was precipitated by adding 200 µL of 1 M Tris-HCl pH 8 and 1 ml of pure ethanol, followed by overnight incubation at -80°C. DNA fragments were recovered by centrifugation for 30 min at 20,000 g and 4°C. The resulting pellets were resuspended in 20 µL of TE (10 mM Tris-HCl pH 8, 1 mM EDTA) and resolved on a 4% agarose gel in 0.5X TAE buffer (20 mM Tris, 10 mM acetic acid, 0.5 mM EDTA) for 2 h at 4 V/cm. The resulting bands for each of the three biological replicates were gel eluted and 15 ng of DNA was end-repaired using the NEBNext Ultra II End Prep mix (NEB) by incubating for 30 min at 20 °C and 30 min at 65 °C. The DNA was subsequently purified with 1.8 x volume AMPure beads (Beckman Coulter), followed by ligation with Illumina TruSeq adapters (15 nM in a 25 µL reaction) and 2 µL Quick T4 DNA ligase (NEB) for 15 min at 20 °C. Then another round of AMPure beads purification was done (1.2x ratio) followed by PCR amplification using primers corresponding to the i5 and i7 extremities of the adapters. Libraries were sequenced on an Illumina NextSeq500 instrument using NextSeq 500/550 High Output Kit v2 (300 cycles) and a Paired-End 2 × 75 bp run. Demultiplexing was performed with bcl2fastq2 v2.18.12. Adapters were trimmed with Cutadapt v1.15, and only reads longer than 10 bp (between 70 and 80 million reads per sample) were kept for further analysis.

### Total RNA isolation and sequencing

To prepare samples for RNA sequencing, 25 ml of Ravot medium was inoculated at 1/100 dilution with fresh *T. kodakarensis* preculture. Total RNA was extracted from 20 ml of exponentially growing cultures (6 h of culture, approximately 2 × 10^9^ cells/ml) using a NucleoSpin RNA set for NucleoZOL (Macherey Nagel). DNA was eliminated from samples using a TURBO DNA-free kit following the manufacturers protocol (Ambion). For both the ΔRG and the WT strain biological replicates were prepared from four independent cultures. Total RNA quality and concentration was assessed using an Agilent 2100 Bioanalyser RNA 6000 Nano assay. Ribosomal RNA depletion was performed using siTOOLs riboPOOLs module, and NGS RNA-Seq libraries made using the NEBNext® Ultra™ II Directional RNA Library Prep Kit for Illumina® according to manufacturer’s instructions. Library size and adaptor contamination was assessed using the Agilent 2100 Bioanalyser High Sensitivity DNA assay, and DNA concentrations measured with the Qubit dsDNA High Sensitivity assay. For each sample, a minimum of 3 million Paired End 150 bp reads were generated on an Illumina Miseq with unique dual 8 bp indexing.

### TMP-seq and MNase-seq analyses

Reads were first aligned to the full genome of *T. kodakarensis* using the Bowtie2 aligner. Mapped reads were deduplicated, sorted, and indexed with Samtools^40^ and Picard (http://broadinstitute.github.io/picard). Coverage files (BigWig) for CL and noCL samples were generated using the bamCoverage command from the deepTools package^41^. Profile subtraction and log2 ratio computation were performed using the bigwigCompare command from the same package. Averaged BigWigs were generated using the bigwigAverage command. Heatmaps and average read distributions at genomic regions were plotted using the computeMatrix, plotHeatmap and plotProfile commands. Z-normalization, inverse hyperbolic sine transformation and smoothing of the BigWigs files were performed using the GenomicRanges package in R. Crosslinking level (CLL) was calculated using a hyperbolic sine transformation (arcsinh) of the crosslinked/non-crosslinked DNA ratio, instead of a log2 ratio, to account for negative z-scores. Correlation matrices were calculated using the multibigwig command and plotted using ggplot2 in R. Loess smoothing of average profiles was performed using the ggplot2 package. Scatter plots were generated with matplotlib in Python.

### RNA-seq analysis

Adapters were trimmed with Cutadapt and only reads longer than 10 bp were kept for further analysis. Reads were first aligned to the rRNA genes of *T. kodakarensis* using Bowtie2. Mapped reads were discarded and only unmapped reads were then aligned to the full genome of *T. kodakarensis* and analysed. Samtools was used to index and sort reads. The number of reads per feature was counted using HTseq-count. Differential analysis was performed using DEseq2. Coverage was calculated using the Deeptools bamCoverage command. Transcription units were predicted in silico using Operon mapper^42^.

## RESULTS

### Increased cell size and impaired growth in *T. kodakarensis* cells that lack reverse gyrase

To assess the importance of RG for the growth of *T. kodakarensis*, we constructed the ΔRG strain, which carries a markerless deletion of the *rgy* gene, and compared its growth to the WT strain at three different temperatures: 85°C (the optimal growth temperature) and two supra-optimal temperatures, 90°C and 93°C. Doubling time of ΔRG cells grown in rich medium increased approximately threefold during the exponential phase at both 85°C and 90°C. No growth was observed for the ΔRG strain at 93°C, whereas the WT strain was still able to proliferate at this temperature (**Figure 1A, Figure S1**), in agreement with previous work^13^.

**Figure 1.**
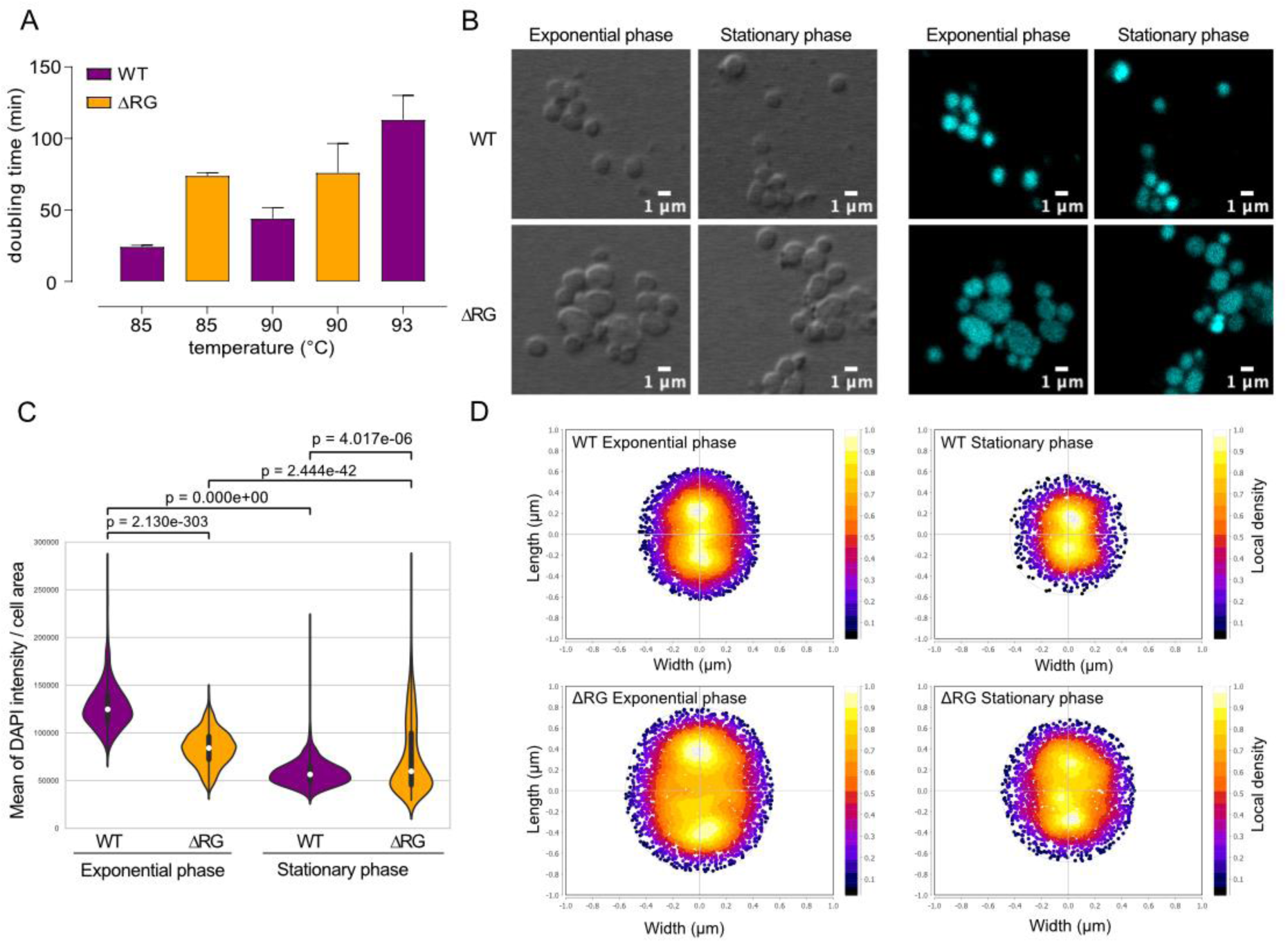
The impact of reverse gyrase deletion on *T. kodakarensis* growth and nucleoid dimensions. A. Mean doubling time of WT and ΔRG cells in exponential growth phase for indicated temperatures. The error bars correspond to standard deviation from the mean of three independent experiments. B. Confocal microscopy images of WT and ΔRG cells in exponential or stationary growth phase. The DNA was stained with DAPI and DIC (left panels) and fluorescence (right panels) images were collected. C. Violin plot showing the distribution of the mean fluorescence signal per cell normalised by the area of the cell. The mean value and first and third quartiles are depicted by a white dot and thick line, respectively. The distributions are significantly different for all conditions, as determined by Mann-Whitney U tests (two-sided) and the corresponding p-values are indicated at the top of each plot. Number of cells per conditions: WT exponential phase: 1138, WT stationary phase: 1096, ΔRG exponential phase: 1164, ΔRG stationary phase: 1208. D. Density plots showing average size and shape of WT and ΔRG cells in exponential or stationary phase. The fluorescence maxima density scale is shown to the right of each plot.

Confocal microscopy of DAPI-stained cells in exponential or stationary phase revealed that the DNA occupied the entire volume of the coccoid cells, in both WT and ΔRG cells, with fluorescence being most intense at the centre of the cells (**Figure 1B-D**). Both WT and ΔRG stationary-phase cells were, on average, smaller and exhibited lower fluorescence intensity per cell area compared to exponentially growing cells (**Figure 1C, Figure S2**). Notably, exponentially growing ΔRG cells were, on average, larger (62% increase in cell area) and showed a significantly lower mean fluorescence intensity, a parameter that can be used as an indirect measure of cellular DNA concentration (**Figure 1D, Figure S2**).

Collectively, these data show that RG is not essential at the optimal growth temperature of *T. kodakarensis*, but its absence results in a markedly reduced growth rate and increased cell size. Confocal microscopy further revealed that, in these enlarged cells, the DNA filled almost the entire cell volume, suggesting that DNA content, distribution and compaction might be altered in the ΔRG strain.

### Widespread deregulation of gene expression and activation of the stress response in reverse gyrase-deficient *T. kodakarensis*

We next performed RNA-seq to assess how the loss of RG affected gene expression. Differential gene expression analysis revealed that a large proportion of genes, 1323 out of 2289 protein coding genes, were significantly deregulated (fold change (FC) > 1.25, adjusted p-value < 0.01) (**Figure S3A**), including 316 genes that were deregulated by more than two-fold. Differentially expressed genes (DEGs) were evenly split between up-regulated (660 genes) and down-regulated (663 genes). A small subset of DEGs (54 in total) exhibited strong expression changes (FC > 4) and those corresponded to stress-related genes such as flagella and genes in chemotaxis operons (upregulated) as well as iron uptake and membrane transport proteins (downregulated). Gene ontology analysis further suggested that deletion of RG negatively affected protein synthesis (COG J and E) and cell division (COG D), while promoting the expression of genes involved in signal transduction and motility (COG N and T) (**Figure S3C**).

Beyond this general transcriptional response, we specifically investigated whether the loss of RG affected the expression of DNA topoisomerases, nucleoid associated proteins (NAPs), histones or genes involved in DNA repair (**Figure 2E, Table S2**). Among the two housekeeping topoisomerases, Topo III, another type 1A topoisomerase, was mildly upregulated (FC = 1.3), while the expression of Topo VI remained unchanged (**Table S2**). TrmBL2 was the only abundant NAP whose expression was altered (FC = -1.6), whereas the expression of both histones, HtKA and HTkB, remained unchanged. Within DNA repair pathways, the essential homologous recombination pathway, Rad50/Mre11/NurA, in charge of repairing double-strand breaks in DNA, was downregulated (up to FC = -3.2). In the nucleotide excision repair pathway, Hef, which has a critical function in dealing with DNA interstrand crosslinks, was upregulated (FC = 1.3), while Xpb was downregulated (FC = -0.63). Additionally, in the base excision repair pathway, 8-oxoguanine DNA glycosylase, which initiates the repair of the oxidative DNA damage product 8-oxoguanine, was upregulated (FC = 2.9) (**Figure 2E**). Finally, the operon encoding the three Replication Protein A (RPA) paralogs, which bind to ssDNA and protect it from damage during DNA replication and repair, was strongly upregulated (FC = 3.8) (**Figure 2E**).

**Figure 2.**
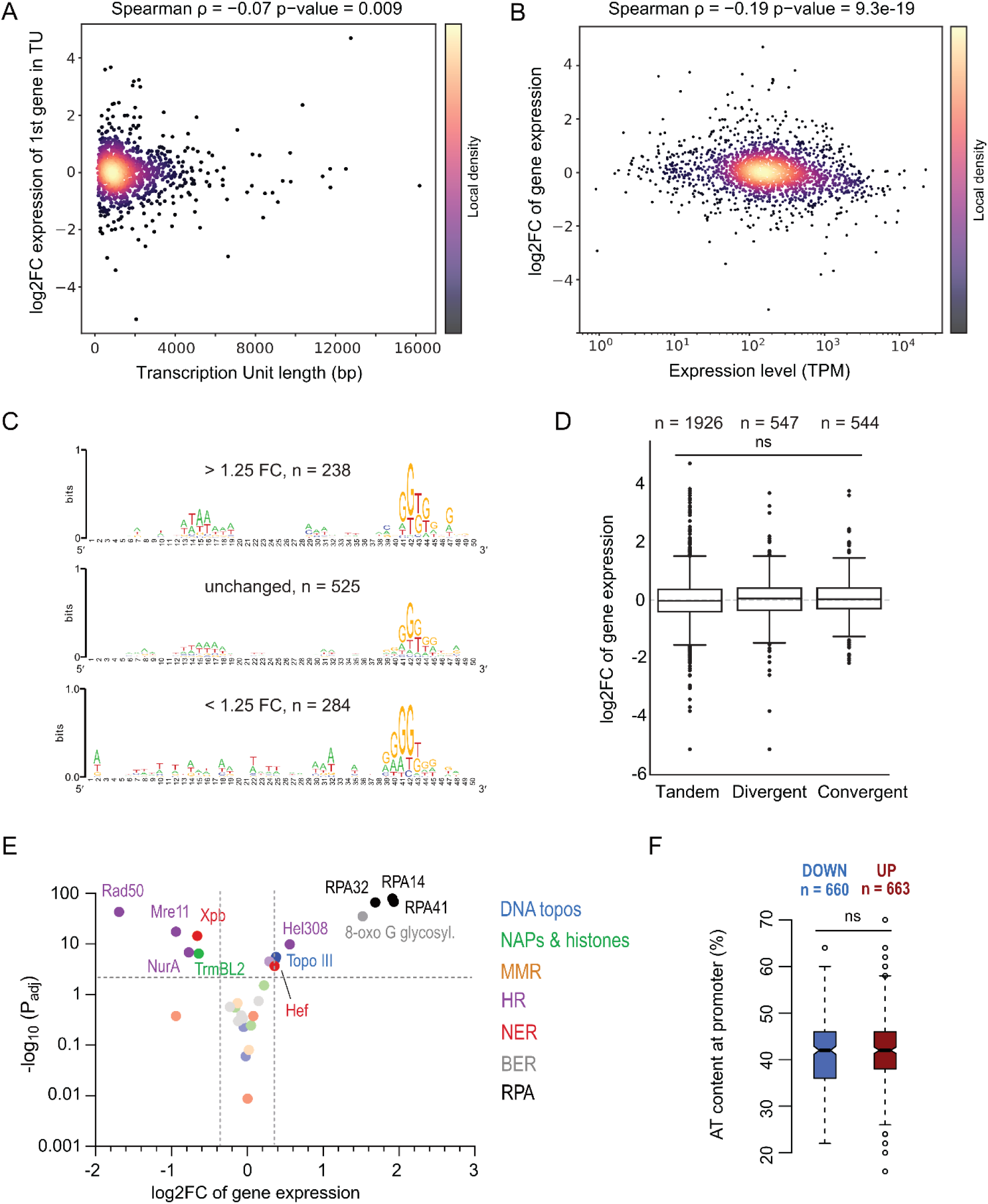
Impact of deletion of reverse gyrase on gene expression in *T. kodakarensis*. A. Scatter plot showing the log2FC of expression of the first gene in a transcription unit (TU) as a function of the length of the transcription unit. B. Scatter plot showing the log2FC of gene expression as a function of the gene expression level in the WT strain. TPM: transcripts per million. C. Sequence logo of promoters classified according to the fold change in gene expression. Only the primary promoter per transcription unit was considered. The number of analysed promoters per category is indicated above each plot. D. Distribution of log2FC values for the three categories of mutual gene orientations: tandem, divergent and convergent genes. In some cases, a gene can be assigned to different categories at the same time. The box shows the median value and interquartile range (25^th^ percentile to 75^th^ percentile) while the individual data points correspond to outliers. The difference in median values between different categories are not statistically significant as measured by Mann- Whitney U tests. E. Differential expression of genes encoding DNA topoisomerases, histones, NAPs and DNA repair proteins. Dotted lines indicate the threshold values for adjusted P value (P_adj_ < 0.01) and fold change (Log2FC ≥ |0.33|). MMR – mismatch repair; HR – homologous recombination; NER – nucleotide excision repair; BER – base excision repair; RPA – Replication Protein A. F. Distribution of AT content values for promoters of downregulated or upregulated genes. The number of genes in each category is indicated on the top of the plot. The difference in median values between the two categories is not significant as determined by Welsh’s two-sample t-test.

To explore the mechanistic basis of this widespread deregulation, we analysed the relationship between fold change in expression of the first gene in transcription unit (TU), representative of the expression of the whole operon, and TU length (**Figure 2A)**, gene expression level (**Figure 2B**), genic AT content (**Figure S3D**), mutual gene orientation **(Figure 2D, Figure S3E**), promoter sequence (**Figure 2C)** and promoter AT content **(Figure 2F, Figure S3B**). Among these variables, only the TU length and, to a greater extent, gene expression level, were significantly negatively correlated with the fold change in gene expression, indicating that upon RG deletion long and highly expressed genes tended to be less expressed while short and lowly expressed genes tended to be more expressed.

Taken together, these data reveal 1) deregulation of DNA repair pathways responsible for resolving oxidative and strand break lesions in the ΔRG strain, suggesting that the cells experience DNA damage commonly induced by high temperature, 2) a global reduction in the dynamic range of transcription in the ΔRG strain, characterized by a broad modulation of transcriptional output, in which lowly expressed genes are upregulated whereas highly expressed are downregulated. These transcriptional changes may arise from alteration in the chromatin structure or redistribution of DNA supercoiling across the genome. To test this, we assessed both parameters genome-wide.

### Histone occupancy remains stable in the absence of reverse gyrase

The genome of *T. kodakarensis* is densely coated with histones that form tetramers on DNA and can further assemble into variable-length oligomers through the stepwise addition of histone dimers, progressively wrapping more DNA^43,44^. To determine whether this chromatin structure is altered in the absence of RG, we assessed histone occupancy across the genomes of WT and ΔRG strains using micrococcal nuclease digestion of chromatin coupled with deep sequencing (MNase-seq).

In both strains, the MNase footprints of histone complexes exhibited a typical ladder-like pattern with 60, 90, 120 and 150 bp bands (**Figure 3A**) corresponding to DNA wrapped by two, three, four and five histone dimers, respectively. As measured from this profile, the stoichiometry of bands within each sample was not significantly different between the two strains (**Figure 3B)** suggesting that the capacity to form histone oligomers was not impaired by RG deletion.

**Figure 3.**
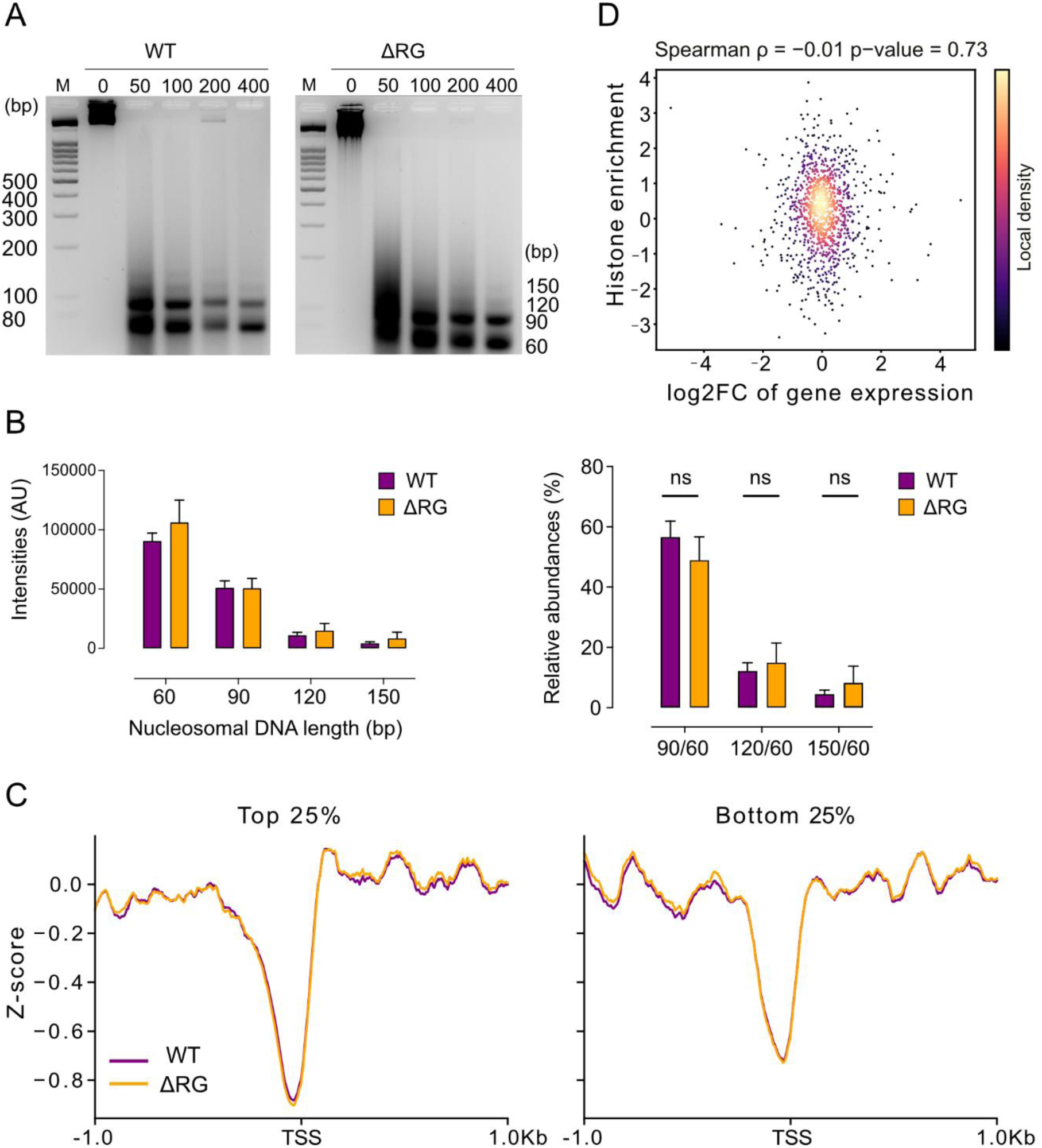
Histone occupancy remains unchanged in the absence of reverse gyrase. A. Footprints of histone complexes isolated from the WT (left panel) or ΔRG (right panel) strain using MNase digestion. The sample digested with 200 U of MNase was used for deep sequencing. B. Histone oligomer abundance. The bands corresponding to DNA fragments protected from MNase digestion were quantified by densitometry from three independent replicates (left panel) and the density ratio of each band to the major 60 bp band was calculated (right panel). The ratios are not significantly different (ns) between WT and ΔRG strains. C. Average histone coverage across genes centred on the transcription start site (TSS) in the WT and ΔRG strains. Read coverage was transformed to z-scores (mean = 0, SD = 1) per replica and averaged. The genes were stratified into top and bottom 25% expressed genes and analysed separately. D. Histone occupancy enrichment at promoters (log2 ratio of histone occupancy in ΔRG versus WT) versus fold change in gene expression. The two variables are not correlated (Spearman test P=0.73).

Deep sequencing of the bands revealed highly similar histone coverage distributions across replicates (n=3) and between the two strains (Spearman correlation > 0.9) (**Figure S4A**). Plotting the aggregate signal distribution across all predicted transcription units (TUs) revealed low histone occupancy in the promoters immediately upstream of transcription start sites (TSS) and downstream of predicted transcription end sites (TES) (**Figure S4B**), consistent with previous reports^43,45^. As the genome of *T. kodakarensis* (as in all prokaryotes) is tightly packed with protein-coding genes^46^ and most intergenic regions are <100 bp, with many <50 bp, we cannot exclude some overlap between promoter and terminator sequences of adjacent TUs. Further comparison between genes belonging to the highest and lowest quartile transcription levels again showed nearly identical profiles in both strains (**Figure 3C**). Finally, histone enrichment across promoters (log2 ratio of ΔRG vs WT strain) (**Figure 3D**) or across core promoters (limited to 50 bp upstream from TSS) (**Figure S4C**) was not correlated with fold change in gene expression.

These findings suggest that despite widespread transcriptional changes, histone occupancy at gene-regulatory regions remained unaffected in the RG mutant, indicating that the loss of RG in *T. kodakarensis* influences gene expression through mechanisms independent of histone associated chromatin accessibility.

### Psoralen photobinding detects chromosomal supercoiling under native growth conditions of *T. kodakarensis*

Given that RG exhibits positive supercoiling activity *in vitro*, we next investigated whether it affects chromosomal DNA supercoiling using a trimethylpsoralen photobinding assay coupled with deep sequencing (TMP-seq) (**Figure 4A**).

**Figure 4.**
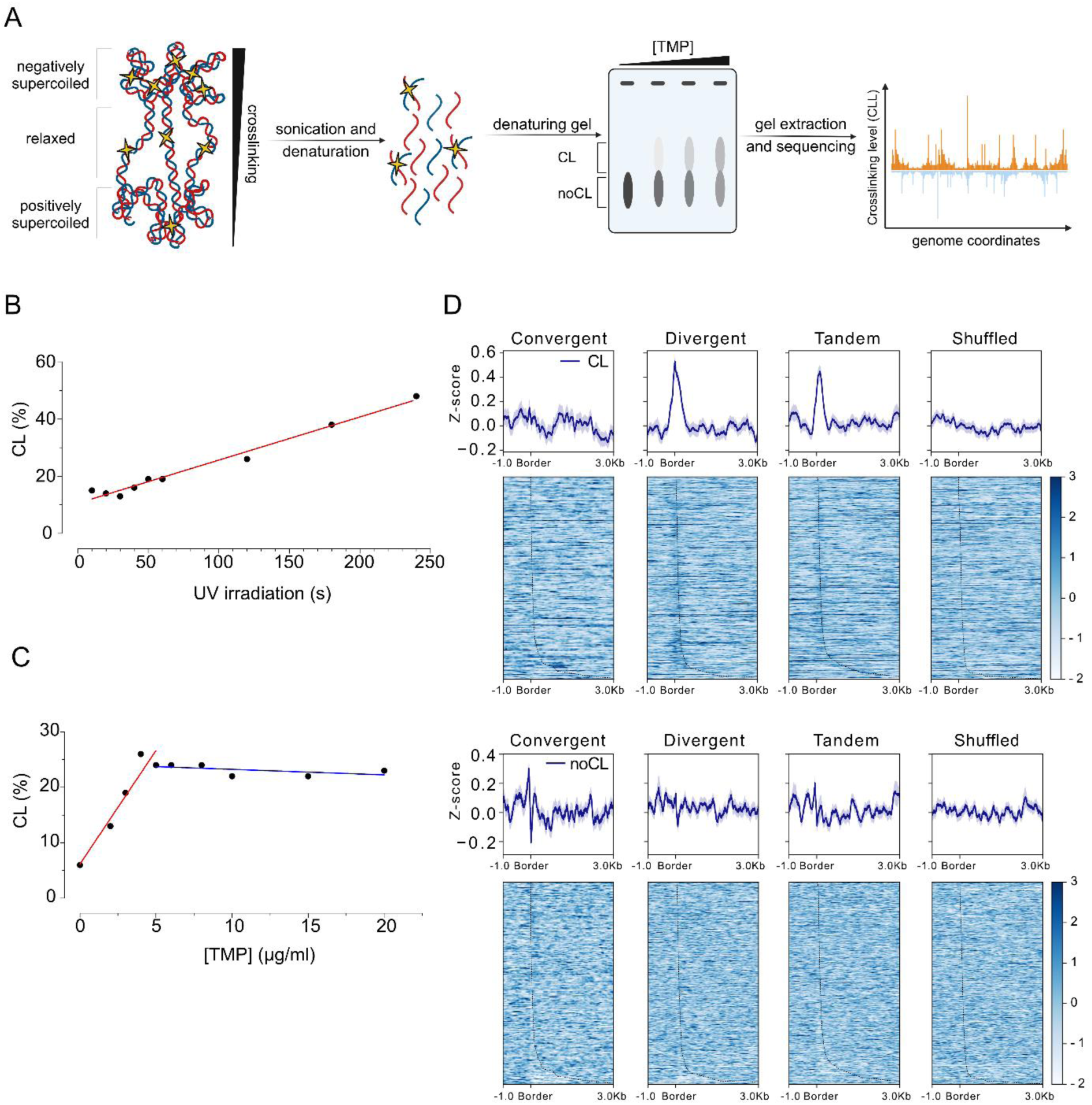
Psoralen photobinding detects DNA supercoiling in *T. kodakarensis*. A. TMP-seq procedure. TMP (yellow stars) preferentially binds to underwound (negatively supercoiled) DNA and forms UV-induced inter-strand crosslinks (CL) with DNA. The DNA is isolated from cells, fragmented by sonication and separated on a denaturing agarose gel. CL DNA migrates more slowly compared to DNA that did not incorporate psoralen (noCL). The CL and noCL DNA fragments are gel-extracted and sequenced separately. Cross Linking level (CLL), calculated as the log_2_ ratio (CL/noCL), provides a continuous picture of psoralen binding as a function of genome coordinate. Created with BioRender.com. B. Effect of UV dose on the formation of TMP-DNA crosslinks. Cells were incubated with 5 µg/ml of TMP and then UV-irradiated (365 nm) at 9.6 kJ.m^-2^.min^-1^ for the indicated amount of time. Crosslinked DNA and non-crosslinked DNA were separated using denaturing gel electrophoresis and quantified using Fiji software (see **Figure S5A**). C. Effect of TMP concentration on formation of TMP-DNA crosslinks. Cells were incubated with varying concentrations of TMP and UV-irradiated for 1 min at 9.6 kJ.m^-2^.min^-1^. Crosslinked DNA and non-crosslinked DNA were separated using denaturing gel electrophoresis and quantified using Fiji software (see **Figure S5B**). D. Distribution of CL (top panel) and noCL (bottom panel) sequencing reads across convergent (n=205), divergent (n=334) and tandem (n=420) TUs. The plots’ reference point is the border of the intergenic region while the dotted line going through the heatmap represents the intergenic region length. Shuffled regions (n=500) with a length distribution matching that of intergenic regions and randomly placed across the genome were used as a control (rightmost column). Read coverage was transformed to z-scores (mean = 0, SD = 1) per replica and averaged. The heat maps show relative levels of TMP incorporation for each TU in a given group (one per row). The bar on the right of the heat maps color-codes the signal intensity.

Since the use of this assay had not previously been reported for hyperthermophiles or archaea, we first tested whether TMP could penetrate *T. kodakarensis* cells. We incubated exponentially growing cells with a fixed TMP concentration and increasing UV doses at 85°C, the optimal growth temperature for *T. kodakarensis*, at which the loss of RG induces minimal DNA damage. This confirmed that TMP photobinding can be established in *T. kodakarensis* (**Figure 4B, Figure S5A)**. The fraction of interstrand TMP-crosslinked DNA increased linearly over a wide range of UV doses (from 1.6 kJ.m^-2^ to 38.4 kJ.m^-2^).

Next, we titrated TMP at a fixed UV dose (28.8 kJ.m^-2^) and found that the crosslinked DNA fraction increased linearly up to 5 µg/ml of TMP before stagnating at higher TMP concentrations (**Figure 4C, Figure S5B)**. Based on these data, we selected non-saturating, low-hit crosslinking conditions (1 µg/ml of TMP and 28.8 kJ.m^-2^ of UV irradiation), corresponding to approximately one crosslink for every 1 – 1.3 kbp of DNA. Under these conditions, 15-19% of DNA was crosslinked and this was consistent across three biological replicates in both the WT and the ΔRG strain (**Figure S5C**). Crosslinked (CL) and non-crosslinked (noCL) DNA were then isolated, used to prepare sequencing libraries, and sequenced.

The distribution profiles of reads were highly similar across replicates (Spearman correlations > 0.85) (**Figure S5D**) and replicates were thus averaged to increase the signal-to-noise ratio.

We analyzed CL and noCL signals based on transcription unit (TU) orientation to assess whether TMP-seq detected supercoiling (**Figure 4D**). According to the twin-domain model^27,47,48^, transcription generates negative supercoiling upstream and positive supercoiling downstream of the elongating RNA polymerase. We therefore expected high levels of negative supercoiling and high levels of TMP binding in intergenic regions (e.g., promoters) of divergently transcribed genes and high levels of positive supercoiling and low levels of TMP binding downstream the termination sites of convergently transcribed genes. Consistent with these predictions, no TMP enrichment was observed in intergenic regions of convergent TUs or in shuffled genomic controls. In contrast, the CL dataset exhibited a clear TMP enrichment at intergenic regions of divergent TUs. TMP was also, on average, enriched in intergenic regions of tandem TUs. In the codirectional TU configuration, the amplitude of the torsion signal is lower than in divergent-TU configuration, indicating that some of positive torsion and negative torsion may cancel out in those regions. The noCL reads showed no bias related to TU orientation, as expected, since noCL DNA lacks topology-sensitive interstrand TMP crosslinks (**Figure 4D**).

Overall, these results establish TMP-seq as a robust method for probing chromosomal DNA supercoiling in *T. kodakarensis*. The agreement between TMP enrichment patterns and the twin-domain model of transcription suggests that this model can be extended to the archaeal domain of life.

### Loss of reverse gyrase affects supercoiling at regulatory regions in the genome

The cross-linking level (CLL), defined as the ratio of CL to noCL signals, provides a quantitative measure of psoralen intercalation across the genome. We analyzed CLL at the scale of individual TUs, aligned at TSS and TES, and observed elevated CLL at both positions, with a stronger signal at TSS than TES sites (**Figure 5A**). If negative supercoiling generated in the downstream promoters influences CLL at upstream TESs, then tandem TUs should exhibit increased signal downstream of TES compared to the convergent TUs. Consistently, the heatmaps show that most of the CLL signal downstream of the TES originates from the nearest downstream TSSs of a tandem TU located within 500 bp (**Figure S8**). We therefore interpret the elevated CLL at TES as a consequence of the high gene density and short intergenic distances^46^. Within gene bodies, CLL was reduced compared with TSS, most likely because positive and negative supercoiling generated by pairs of elongating polymerases may cancel out within gene bodies, thereby reducing TMP intercalation.

**Figure 5.**
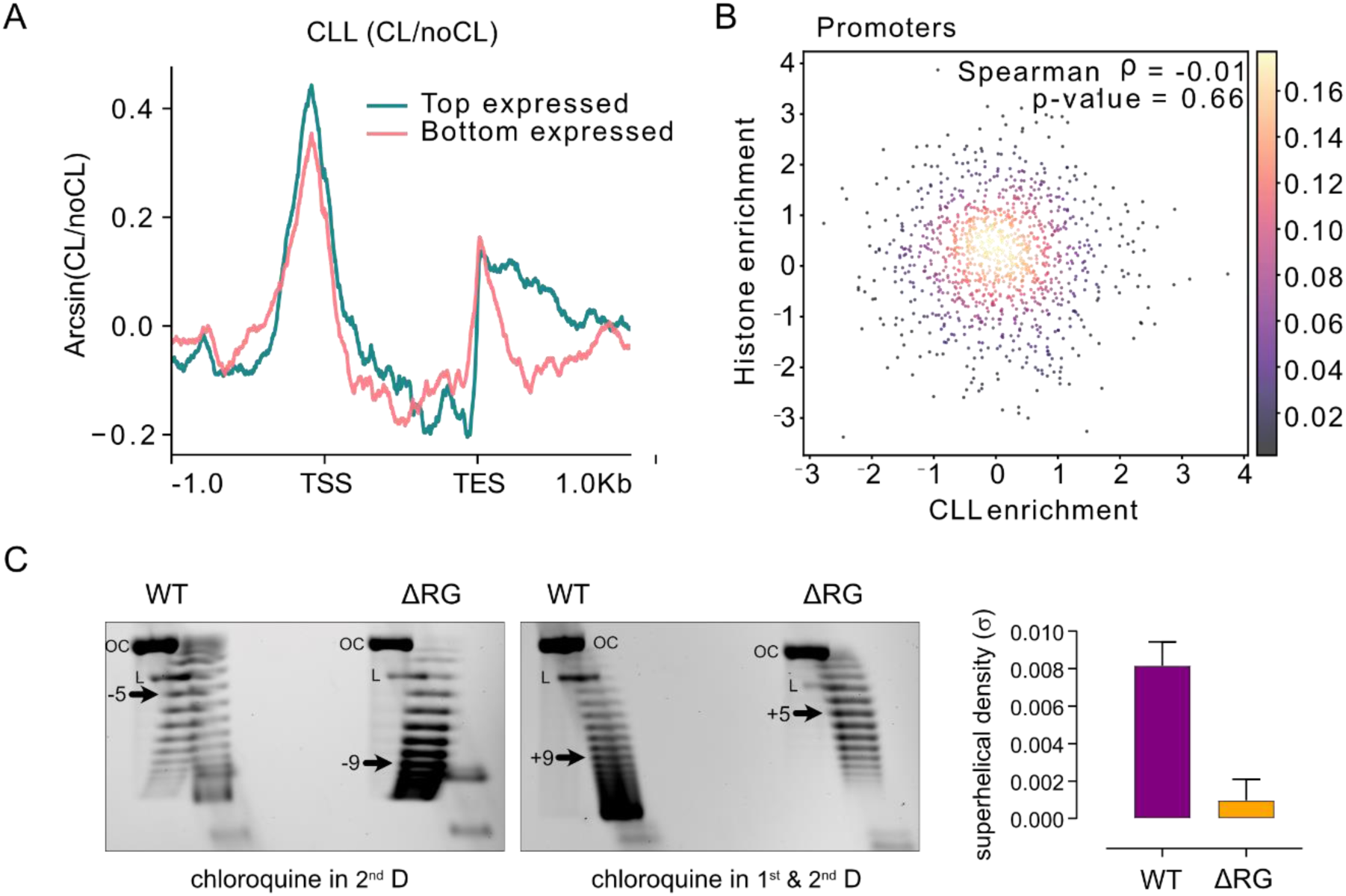
Distribution of DNA supercoiling on the chromosome of *T. kodakarensis*. A. Distribution of WT CLL signal across bottom (n=273) and top (n=272) expressed TUs plotted centred at TSS and TES. Read coverage was z-score normalized per replicate (mean = 0, SD = 1), averaged across replicates, and the resulting signals (CL/noCL) were compared as arcsinh-transformed ratios (CLL). B. Correlation between histone occupancy enrichment (log2 ΔRG/WT) and TMP enrichment (log2 CLL_ΔRG_/CLL_WT_) at promoters of expressed TUs. Each dot corresponds to a single TU. C. Left and middle panel: pTPTK3 plasmid topoisomers were resolved using 2D gel electrophoresis with addition of intercalating agent chloroquine in 2^nd^ dimension (left panel) or both in 1^st^ and 2^nd^ dimension (right panel). Major topoisomers were determined from band density measurements and are indicated with arrows. The numbers correspond to the number of positive or negative supercoils. OC stands for open circular (relaxed) plasmid DNA and L stands for linear form of plasmid DNA. Right panel: the superhelical density (σ) was calculated after correcting for the contribution of chloroquine and temperature. The mean σ value and standard deviation from the mean value were calculated from three independent replicates (**Figure S7**).

To examine the relationship between transcription and supercoiling, we stratified TUs by expression level (top *versus* bottom, based on the ranked WT RNA-seq data). Highly expressed (top) TUs displayed elevated CLL at TSS and downstream of TES relative to lowly expressed (bottom) TUs (**Figure 5A**).

Because CLL enrichment showed no significant correlation with histone occupancy across TSS and TU bodies (Spearman’s ρ = −0.01, **Figure 5B**, **Figure S6A**), these changes in CLL likely reflect alterations in DNA topology rather than changes in chromatin accessibility.

Next, we transformed the ΔRG strain with a replicative plasmid pTPTK3 (4.7 kbp) to test topoisomerase activity of RG *in vivo*. Using 2D gel electrophoresis, we determined the average superhelical density (σ) of the pTPTK3 isolated from exponentially growing cultures of ΔRG and WT strains (**Figure 5C, Figure S7**). While the major topoisomer of pTPTK3 isolated from the WT was positively supercoiled (3-4 positive supercoils), plasmid DNA extracted from the ΔRG strain exhibited a relaxed topology (0-1 positive supercoils) (**Figure 5C**). These data support the notion that RG has positive supercoiling activity *in vivo* and testify that RG activity is not masking a hitherto overlooked negative supercoiling activity in *T. kodakarensis*.

To assess the impact of RG loss on chromosomal DNA supercoiling, we compared CLL profiles between the WT and ΔRG strains. Subtracting WT CLL from ΔRG CLL (ΔCLL), a metric that corrects for chromatin and sequence bias (**Figure S6B-C**), revealed increased values at TSS and TES of TUs in ΔRG (**Figure 6A-B)**. We interpret such increase in ΔCLL as a reduced level of positive supercoiling or a net gain of negative supercoiling following RG loss. We find that highly transcribed TUs accumulate more negative supercoiling in their promoters than weakly transcribed Tus (**Figure 6C**, **Figure S6D**) in ΔRG relative to WT. This redistribution of supercoiling may also account for the transcriptional changes detected by RNA-seq where strongly expressed TUs were downregulated, while weakly expressed TUs were upregulated (**Figure 6D**, **Figure 2B**). Consistent with this hypothesis, the ΔCLL at the most downregulated genes in RG was higher than at non-deregulated genes, whereas the most upregulated genes showed a lower ΔCLL, although this difference was not statistically significant (**Figure 6E**). Notably, in WT cells, the genes that become upregulated or downregulated in ΔRG exhibit, respectively, lower or higher basal expression compared to non-deregulated genes (**Figure 6D**). We therefore propose that RG deletion increases negative supercoiling at highly active promoters thereby hindering transcription initiation^49^, while a modest increase in negative supercoiling at weak promoters may facilitate DNA melting and gene expression activation^49,50^.

**Figure 6.**
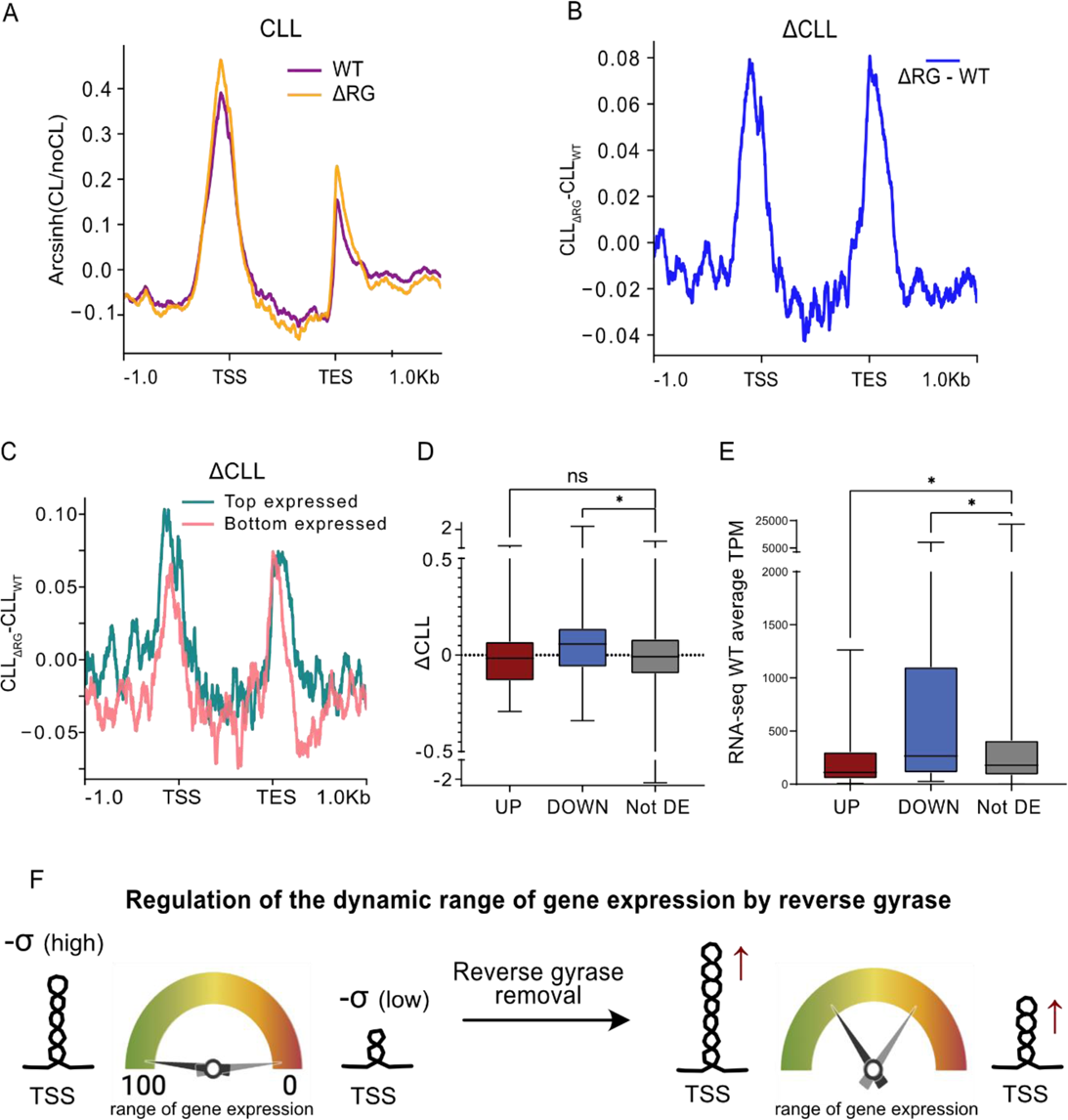
Impact of reverse gyrase loss on chromosomal supercoiling in *T. kodakarensis*. A. Distribution of CLL signal across for all TUs in WT (violet) and ΔRG (orange) strain. B. The difference between WT and ΔRG CLL (ΔCLL) at all TUs. C. ΔCLL across bottom (n=273) and top (n=272) expressed TUs. D. Box-and-whisker plots of ΔCLL per gene, stratified by RNA-seq differential-expression (DE) status (Upregulated n=67, Downregulated n=48, Not DE n=2071). DE genes were defined as *padj* < 0.01 and |log2FC| ≥ 1.5. Points are individual genes. Boxes show median and IQR with whiskers to 1.5×IQR. Upregulated in red, Downregulated in blue, Not DE in gray. Group differences were assessed by Kruskal–Wallis test (up-/downregulated pairwise comparison with Not DE). E. Box-and-whisker plots of gene expression in WT (TPM), displayed and annotated as in panel D. F. Working model: compaction of the dynamic range of gene expression after RG removal. The loss of RG topoisomerase activity leads to increase of supercoiling at promotors whereby highly expressed genes lose output, while lowly expressed genes gain modestly, yielding a narrower transcription signal distribution.

Beside detecting ΔCLL changes at TUs (**Figure 6A-B**), we also identified specific loci across the genome with exceptionally high CLL signals. The proviruses TKV2-TKV4 were among the most prominent, showing elevated CLL in the WT strain and further increases upon RG loss (**Figure S6E**). Notably, the TKV genes are AT-rich and histone depleted. While we cannot exclude that high AT content favours TMP intercalation (**Figure S6B-C**), this result could also suggest that RG may protect genomic regions particularly prone to destabilization/breakage by introducing positive supercoils. Such a mechanism could help prevent detrimental topological changes under high-temperature conditions.

## DISCUSSION

The integrity of DNA in hyperthermophilic organisms is constantly threatened at a magnitude that well-studied mesophilic organisms do not face. To prevent the spontaneous loss of native DNA structure, hyperthermophiles have evolved a number of adaptations such as high intracellular salt concentrations, the presence of polyamines and of small basic proteins that bind duplex DNA (histones and NAPs). However, all of these features are also found in mesophiles^24,51^. The discovery of reverse gyrase (RG), a unique topoisomerase that occurs in all hyperthermophiles but not in mesophiles therefore raised a lot of attention^11,24^. The original idea was that RG, by introducing positive supercoils into DNA, could stabilise DNA structure and prevent strand separation *in vivo*^52^. However, subsequent studies revealed that, 1) both positively and negatively supercoiled plasmid DNA resisted denaturation at very high temperatures^53^ and 2) that hyperthermophilic bacteria have both gyrase and reverse gyrase and contain negatively supercoiled plasmid DNA^54^. Further, we recently showed that it is possible to induce strong negative supercoiling *in vivo* in *T. kodakarensis* without a clear growth defect^33^. All of these findings challenge the model that positive supercoiling is advantageous for hyperthermophiles because it provides genome-wide stabilization of the DNA double helix. In this work, we demonstrate that RG affects DNA supercoiling locally and specifically at promoters, suggesting that its primary role is to act on a subset of genomic regions that constitute hotspots for DNA destabilization and breakage at high temperatures. Notably, a parallel study by Yamaura et al. took different approaches (discussed later) and reached highly concordant conclusions, lending reciprocal support to both studies.

### Loss of RG induces stress response and changes in nucleoid size

Consistent with a previous report^13^, the deletion of RG in *T. kodakarensis* led to a slower growth rate at optimal growth temperature and was lethal at temperatures above 93°C (**Figure 1**). We additionally revealed that the slow-growth phenotype was accompanied by an increase in average cell size. Interestingly, the DNA filled almost the entire cell volume of these enlarged cells suggesting that DNA content, distribution and/or compaction was altered in RG-less strain. This could be explained by a global remodelling of nucleoid size and shape via changes in NAP abundance, which is actively investigated in bacteria as one of the most efficient mechanisms in response to stress^55,56^. Our findings suggest that the absence of RG induces stress, as characterized by strong upregulation of the archaellum operon (the archaeal flagella) and chemotaxis genes. In our conditions, the expression of only one major NAP, TrmBL2, was deregulated in absence of RG (**Figure 2**). TrmBL2 is as abundant as each of the two histones^57^ and it was shown, unlike histones, to form thick fibrous nucleoprotein structures suggesting that it significantly contributes to chromatin structure in *T. kodakarensis*^58^. Hence, this protein could offer a starting point for exploring stress-induced remodelling of nucleoid in *T. kodakarensis*.

### RG maintains global transcriptional balance through control of DNA supercoiling at promoters

Our analysis of transcriptomics data revealed a compression of the dynamic range of gene expression (**Figure 2**, **Figure 6D**), which was also reported in bacterial and mammalian cells, following the inhibition of TopoI and the depletion of the Structural Maintenance of Chromosomes (SMC)-encoding gene, respectively^49,59^. A possible interpretation of our transcriptome data is that, following the removal of RG, highly transcribed genes accumulate an excess of negative supercoiling at their promoters, which will not be balanced by RG positive supercoiling activity thus hindering the expression of strong promoters. The lowly expressed TUs on the other hand will benefit from more negative supercoiling at their promoters favouring promoter opening (**Figure 6F**). This is supported by TMP-seq data showing that CLL increase in ΔRG strain is the highest at the promoters of the top quartile expressed TUs (**Figure 6**). This difference was not correlated to histone occupancy changes (**Figure 3**) suggesting that it can be accounted for by increase in negative DNA supercoiling. The full interpretation is likely more complex, involving other factors that modulate DNA structure such as NAPs and SMC as shown in mammalian cells where perturbation of long-range genomic contacts induced a qualitatively similar transcriptional response^59^. Moreover, we cannot exclude that –although modest– the increased expression of TopoIII (type 1A topoisomerase relaxing underwound DNA) might partially compensate for loss of RG activity at TSS. Of note, whereas in bacteria and eukaryotes the effects of altered supercoiling could only be observed transiently or proved lethal, we showed in a previous study^33^ and in this one that *T. kodakarensis* can tolerate a chronic imbalance in supercoiling, highlighting its exceptional capacity to mitigate the consequences of persistent topological stress.

### Psoralen photobinding distribution in *T. kodakarensis* is consistent with twin supercoiled domain model

The unique combination of bacterial-like (small genomes and genes organised in operons) and eukaryotic-like (several origins of replication, presence of histones) genomic features found in archaea, demonstrates the need for in-depth investigations of chromosome architecture in the different clades of archaea. However, the technical challenges in combination with the fact that most if not all archaeal model species are extremophiles, has so far hindered the investigation of archaeal chromosomal supercoiling. In this study, we demonstrate that the supercoiling-sensitive DNA intercalator TMP can be used to generate high-resolution supercoiling maps in *T. kodakarensis*, (**Figure 4 & 5**) thereby providing the first insights into chromosomal DNA topology in Archaea. Notably, the ΔCLL values we measure are in the same range as the torsion observed in a recent study where the signal was corrected for chromatin and sequence bias^60^, suggesting that our protocol robustly detects changes in supercoiling. The TMP binding pattern in *T. kodakarensis* is consistent with the twin-supercoiled domain model^47^, also reported in eukaryotic and bacterial cells^26,27,29,60,61^ suggesting that this model is the norm across the three domains of life. Supercoiling is confined to approximately 0.5 kb around transcription start and end sites in *T. kodakarensis*, similar to what is observed in eukaryotes, where supercoiling accumulates within 1–2 kb of promoters and terminators^27,60,63^. A TMP-seq approach to map supercoiling across the *Escherichia coli* chromosome yielded an order of magnitude higher values with supercoiling twin-domains generated by RNA polymerase complexes spanning 25 kb in each direction^28,30^. This may suggest that transcription-induced dynamic supercoiling is a short-range genomic force in archaea and eukaryotes but not in bacteria where highly expressed genes strongly affect the topology of tens of genes each, creating highly integrated gene circuits. It is tempting to speculate that this difference is related to the presence of nucleosomes in archaea and eukaryotes that may, through wrapping of DNA, act as insulators and limit the diffusion of supercoils upstream and downstream of promoters.

### RG functions in locally stabilising DNA regions prone to heat-induced destabilisation and damage

The observation that RG modulates the topology of gene regulatory regions lead us to propose that RG has a specialised function in stabilising, by introducing positive supercoils, particularly heat-fragile –AT-rich and nucleosome free– genomic regions such as promoters. Uncontrolled promoter destabilization is particularly important threat for hyperthermophiles that live at extremely high temperatures. It therefore stands to reason that these organisms will evolve specific mechanisms to protect promoters. This is consistent with the finding that the number of genes per transcription unit, co-expressed from a single upstream promoter increases with temperature^57^ while the proportion of the genome dedicated to intergenic regions decreases with temperature in bacteria and archaea^64^. Convergent conclusions were reached in the parallel study of Yamaura et al., which showed by ChIP-seq that RG binds to AT-rich sequences and prevents their thermal denaturation at supra-optimal temperature, as evidenced by the accumulation of the ssDNA-binding protein RPA. Interestingly, in ΔRG cells at supra-optimal temperature, RPA was strongly localised within proviruses TKV2 and TKV3 which we have identified as hotspots of TMP signal and sensitive to RG loss (**Figure S6E**). Collectively, the results from Yamaura et al. and from our study are consistent with local action of the RG at AT-rich loci in *T. kodakarensis* both at optimal and supra-optimal temperatures.

In summary, our study provides novel insights into RG function and establishes an experimental framework for investigating *in vivo* topoisomerase activity in archaea. The possibility to map DNA supercoiling across chromosomal DNA of *T. kodakarensis* offers a valuable entry point for understanding both universal and archaea-specific principles governing the relationship between torsional stress, gene expression, chromatin dynamics, and higher-order chromosome organisation.

## ACKNOWLEDGEMENTS

We thank Hiroki Higashibata (Toyo University) for the generous gift of the DAD and ΔRGDAD *T. kodakarensis* strains.

We are grateful to Fedor Kouzine, Nick Gilbert and Catherine Naughton for their advice and insightful discussions regarding the psoralen photobinding protocol, data analysis and data interpretation.

We acknowledge the High-throughput sequencing facility of I2BC (Centre de Recherche de Gif – http://www.i2bc.paris-saclay.fr/) for its sequencing and bioinformatics expertise with particular thanks to Erwin van Dijk, Yan Jaszczyszyn and Céline Hernandez for their technical advice.

We acknowledge Laurence Game, Ivan Andrew and Max Taylor from the LMS Genomic Facility for the RNA sequencing.

The present work has benefited from Imagerie-Gif core facility supported by I’Agence Nationale de la Recherche (FBI ANR-24-INBS-0005 (BIOGEN); SPS ANR-17-EUR-0007, EUR SPS-GSR).

For part of the TMP-seq and MNase-seq analysis computations and data storage were enabled by resources in projects (NAISS 2023/23-603 and NAISS 2023/22-258), provided by the National Academic Infrastructure for Supercomputing in Sweden (NAISS) and the Swedish National Infrastructure for Computing (SNIC) at UPPMAX partially funded by the Swedish Research Council through grant agreements nos. 2022-06725.

## AUTHOR CONTRIBUTIONS

P.V. designed and performed experiments and analysed sequencing data. V.K developed sequencing data pipelines and performed data analyses. A.H. analysed sequencing data. A.D. and F.L. provided technical support. R.LB performed confocal microscopy data analyses. T.W. provided access to necessary equipment and facilities. T.B. and L.B. designed the study and supervised the work. T.B., P.V., L.B and V.K. wrote the first draft of the manuscript. All authors edited the manuscript and approved the final version.

## DATA AVAILABILITY

The sequencing data have been deposited in ArrayExpress (https://www.ebi.ac.uk/biostudies/arrayexpress) under following accession numbers: E-MTAB-15785 (RNA-seq), E-MTAB-15798 (MNase-sseq) and ), E-MTAB-15799 (TMP-seq).

