## Supplementary data for "Localised activity of reverse gyrase at gene regulatory elements"

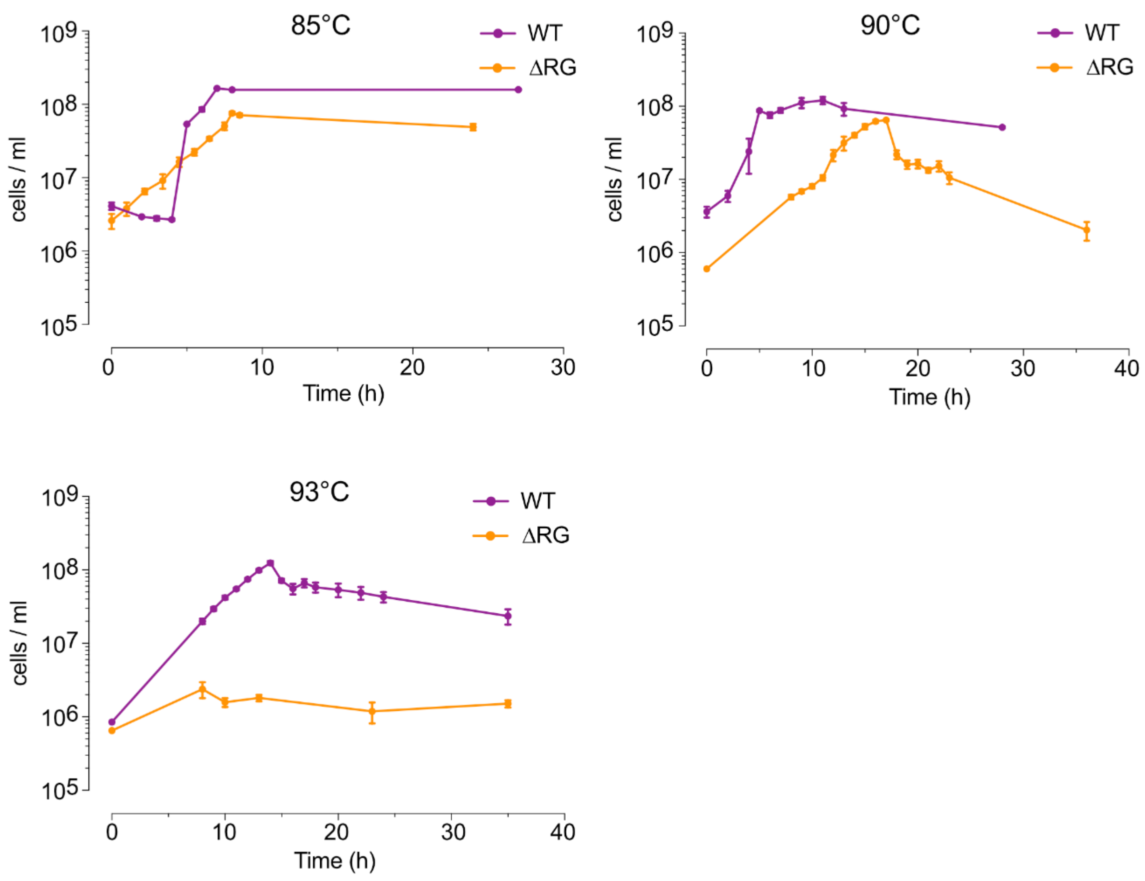


**Figure S1. Growth of *T. kodakarensis* in the absence of reverse gyrase.**

Growth at indicated temperatures was monitored by counting cells using a Thoma chamber. The error bars correspond to standard deviation from the mean value (n = 4).

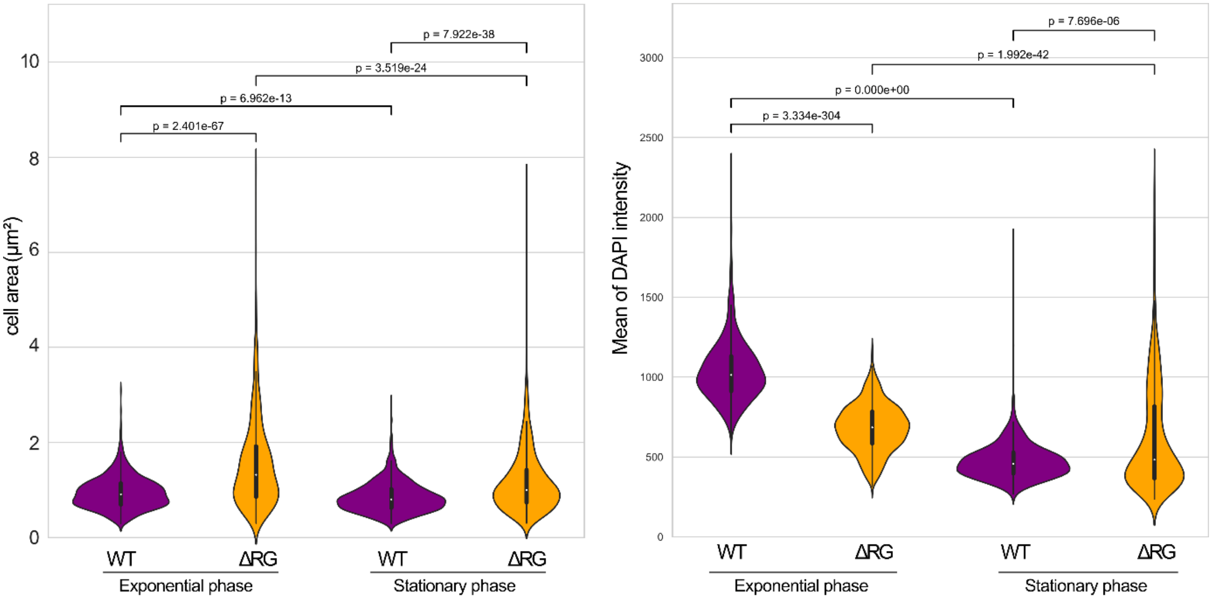


**Figure S2. Measurement of cell area and DNA content in *T. kodakarensis* cells.**

Violin plots showing cell area distribution of *T. kodakarensis* cells in exponential or stationary growth phase (left panel) and DNA content distribution in individual cells measured as mean DAPI florescence intensity (right panel). The mean values for all conditions are significantly different as determined by a Mann-Whitney U test (two-sided). The corresponding p-values are indicated at the top of the plots.

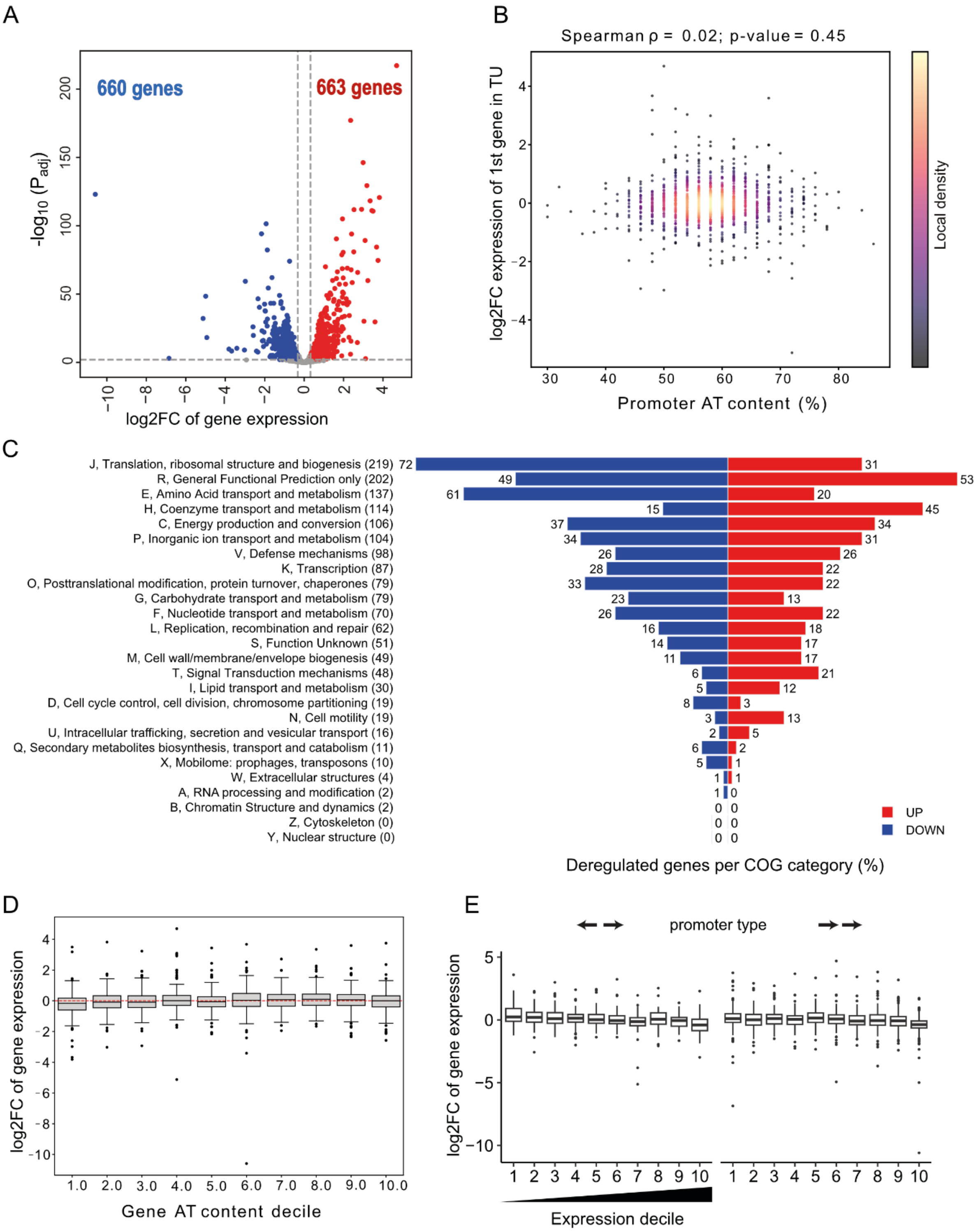


**Figure S3. Global transcriptional response of *T. kodakarensis* in absence of reverse gyrase.**

A. Volcano plot showing the $log_2$ ratio of expression level in the ΔRG strain relative to the wild type. Each dot corresponds to one gene, with blue and red colour indicating downregulated and upregulated genes, respectively. The dotted lines indicate chosen threshold values (adjusted p-value < 0.01 and FC > 1.25).

B. Scatter plot showing the AT content of core promoters (defined as the 50 bp before the ATG of each transcription unit) relative to fold change in expression of the first gene in a transcription unit. The statistical dependence between the two variables was determined using Spearman correlation.

C. Butterfly plot showing the classification of deregulated genes by functional category. COG stands for cluster of orthogolous genes.

D. Box plot showing the fold change in gene expression relative to AT content of genes split by deciles.

E. Box plot showing the fold change in gene expression according to mutual gene orientation split by deciles.

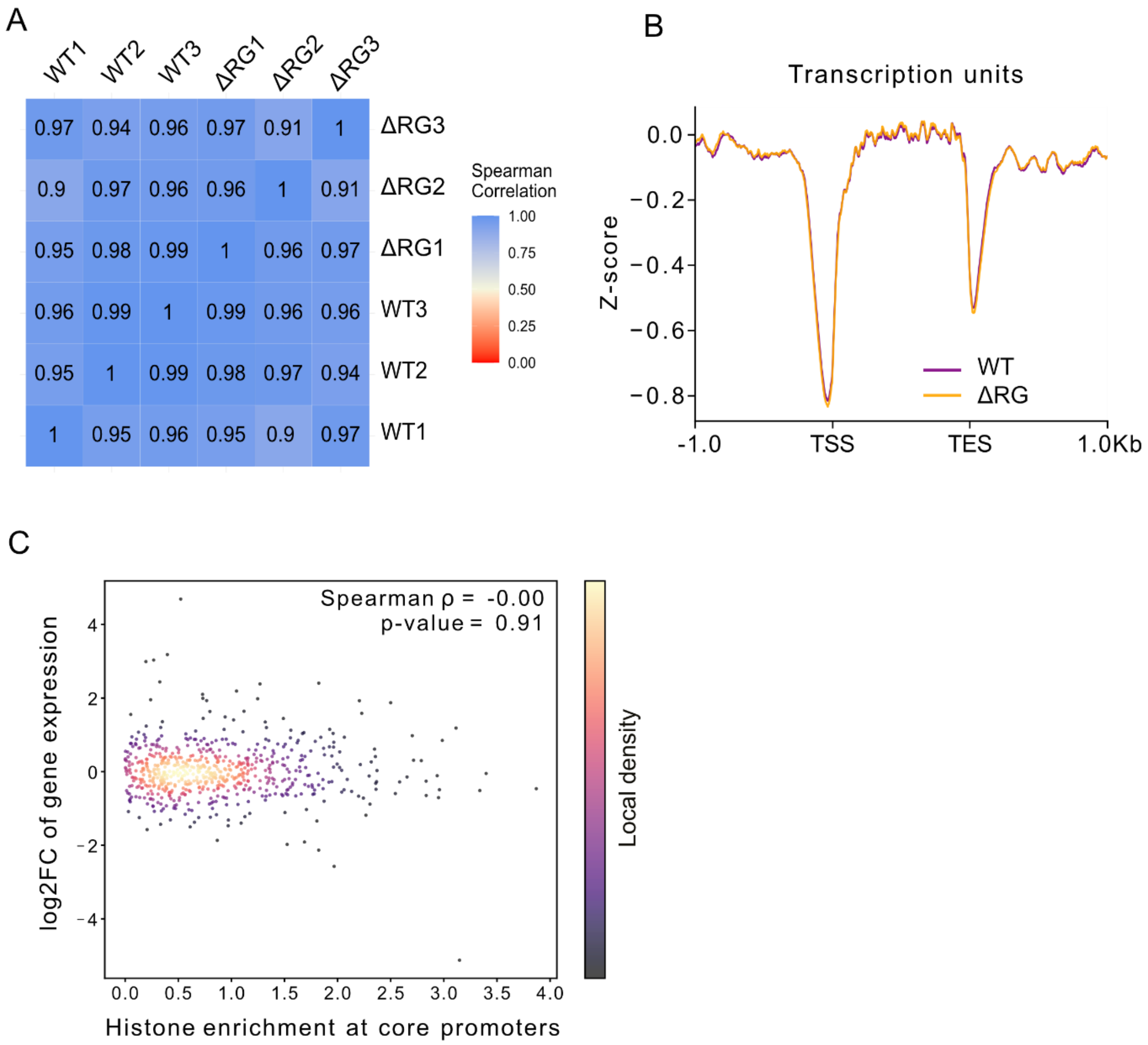


**Figure S4. Analysis of chromatin structure in *T. kodakarensis* using MNase digestion.**

A. Correlation matrix for sequencing read coverage showing the degree of Spearman correlation between replicates (n=3) and the two conditions, the WT and ΔRG strains.

B. Histone occupancy aggregate signal plotted across all identified transcription units centred at the TSS.

C. Scatter plot showing the fold change in gene expression in the ΔRG strain as a function of histone enrichment at core promoters, defined as the 50 bp before the ATG of each transcription unit. The two variables are not correlated as determined by Spearman test. The local density is indicated by the bar to the right of the plot.

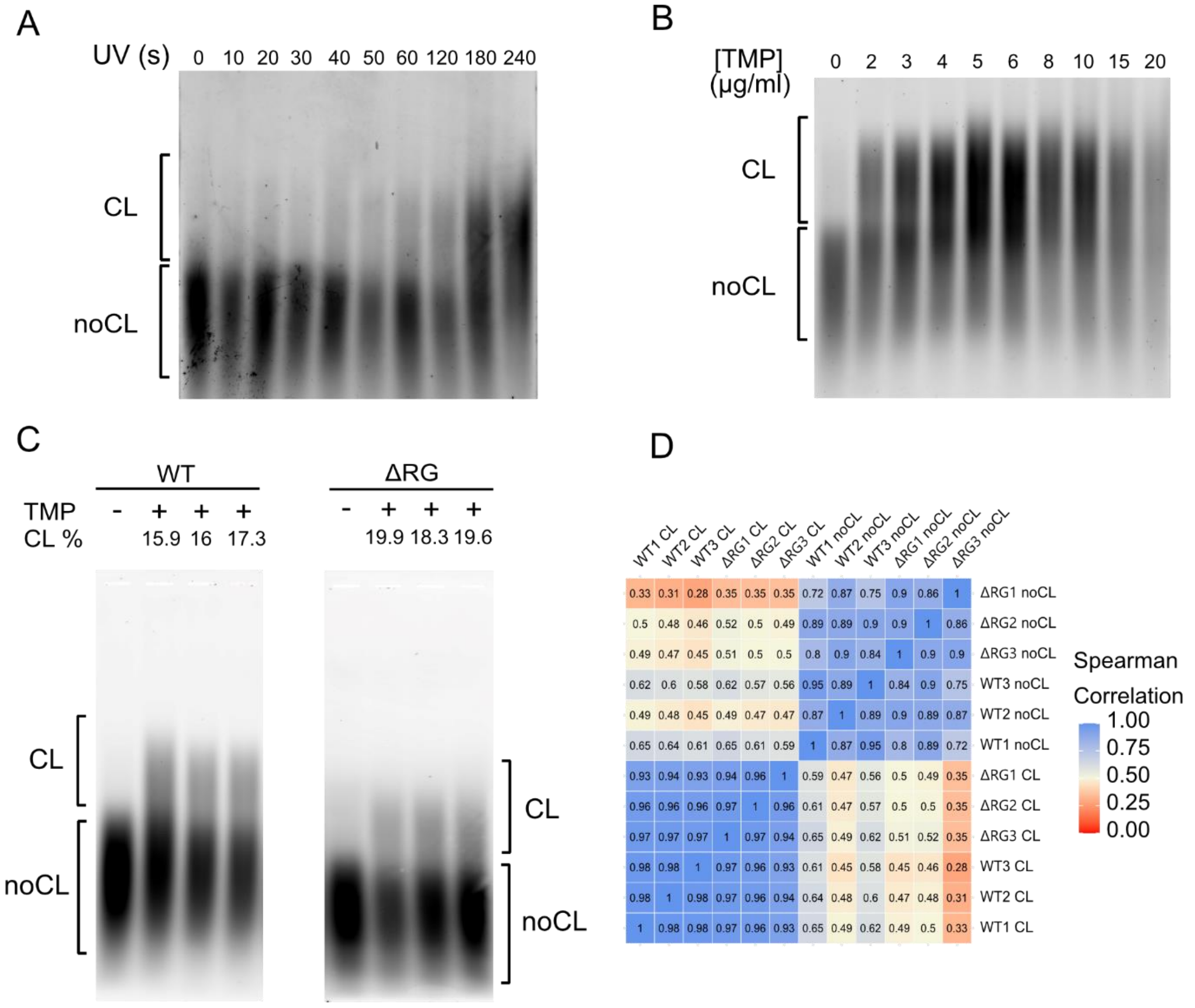


**Figure S5. Establishment of psoralen photobinding assay for *T. kodakarensis*.**

A. Variation of UV dose at fixed concentration of TMP. The cells were UV-irradiated for the indicated amount of time. The genomic DNA was separated under denaturing conditions on an agarose gel. The position in the gel of crosslinked and non-crosslinked fragments is indicated on the side. The crosslink percentage corresponds to the ratio of integrated signal intensity of crosslinked DNA and total DNA per lane.

B. Cultures of *T. kodakarensis* were incubated 10 min with varying concentrations of TMP and UV-irradiated for 1 min (28.8 $kJ.m^{-2}$, 45 W at 4 cm distance). The crosslinked DNA was separated and analysed as for A.

C. DNA samples used for deep sequencing. The ratio CL/total DNA was determined by image analysis. The percentage of CL DNA is indicated above each lane. Control samples received no TMP treatment (-) and for each strain three biological replicates (+) were prepared.

D. Heat map showing Spearman correlation for cross-linked (CL) and non-crosslinked (noCL) datasets for the WT and ΔRG strains.

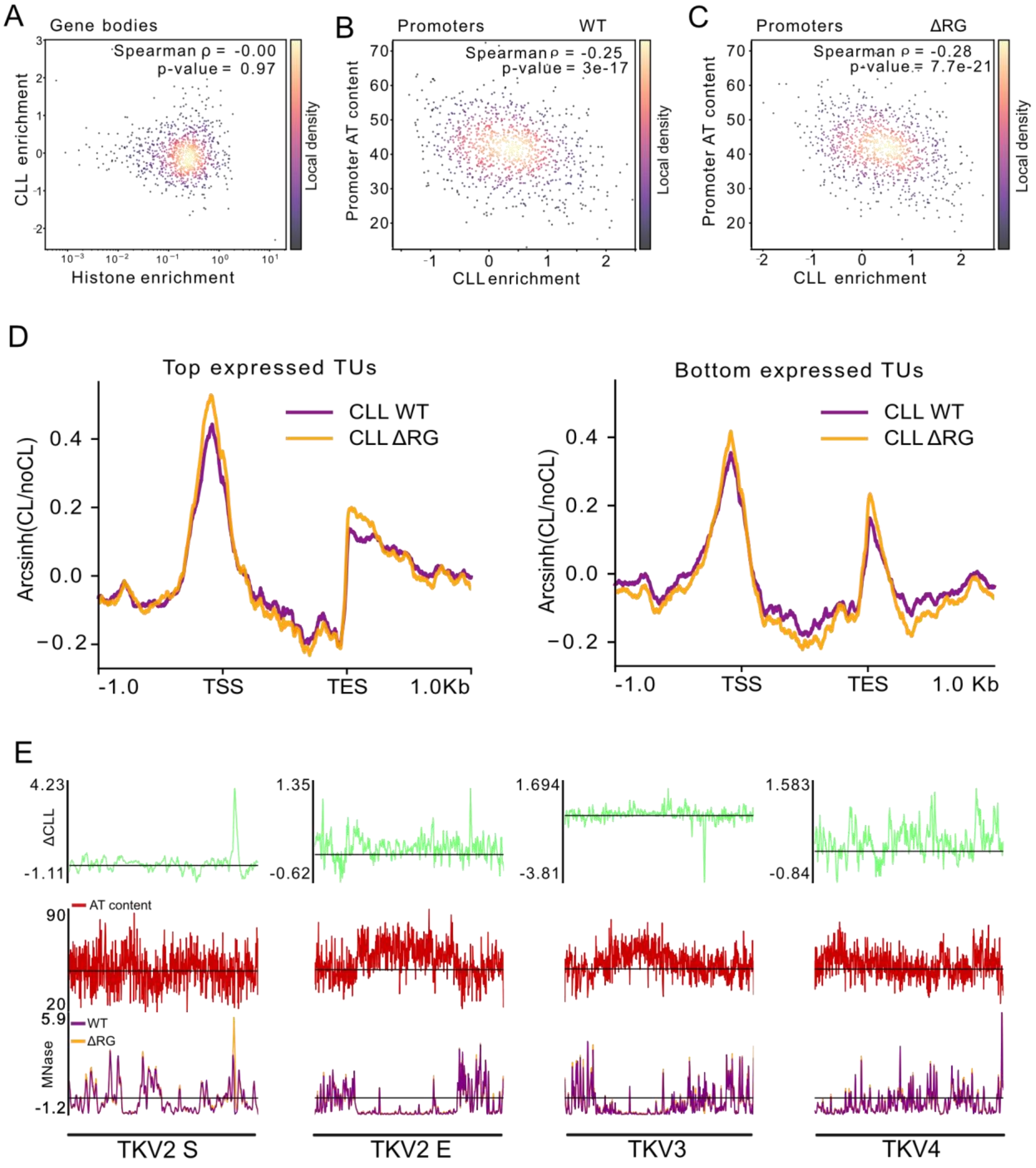


**Figure S6. Supercoiling distribution across the chromosome of *T. kodakarensis*.**

A. Scatter plot showing the CLL enrichment (log2 $CLL_{RG}/CLL_{WT}$) as a function of histone enrichment (log2 RG/WT) across gene bodies. The two variables are not correlated as established by Spearman test. Local density is indicated by the bar to the right of the plot.

B. Scatter plot showing the correlation between AT content at promoters of TUs and CLL enrichment in the WT *T. kodakarensis*. The two variables are significantly correlated.

C. Scatter plot showing the correlation between AT content at promoters of TUs and CLL enrichment in the ΔRG strain. The two variables are significantly correlated.

D. CLL distribution for the WT and the ΔRG strains across bottom (n=273, right panel) and top (n=272, left panel) expressed TUs.

E. Exceptionally high CLL coincide with AT-rich proviral regions in the genome of *T. kodakarensis*. The top plots show the ΔCLL profiles of genomic regions corresponding to three integrated proviral elements TKV2-4 in WT and ΔRG strains. The bottom plots show the distribution of AT content and histone occupancy in these regions. TKV2 spans the genome coordinate boundary; in the browser it is shown as two segments—TKV2 start (S) and TKV2 end (E).

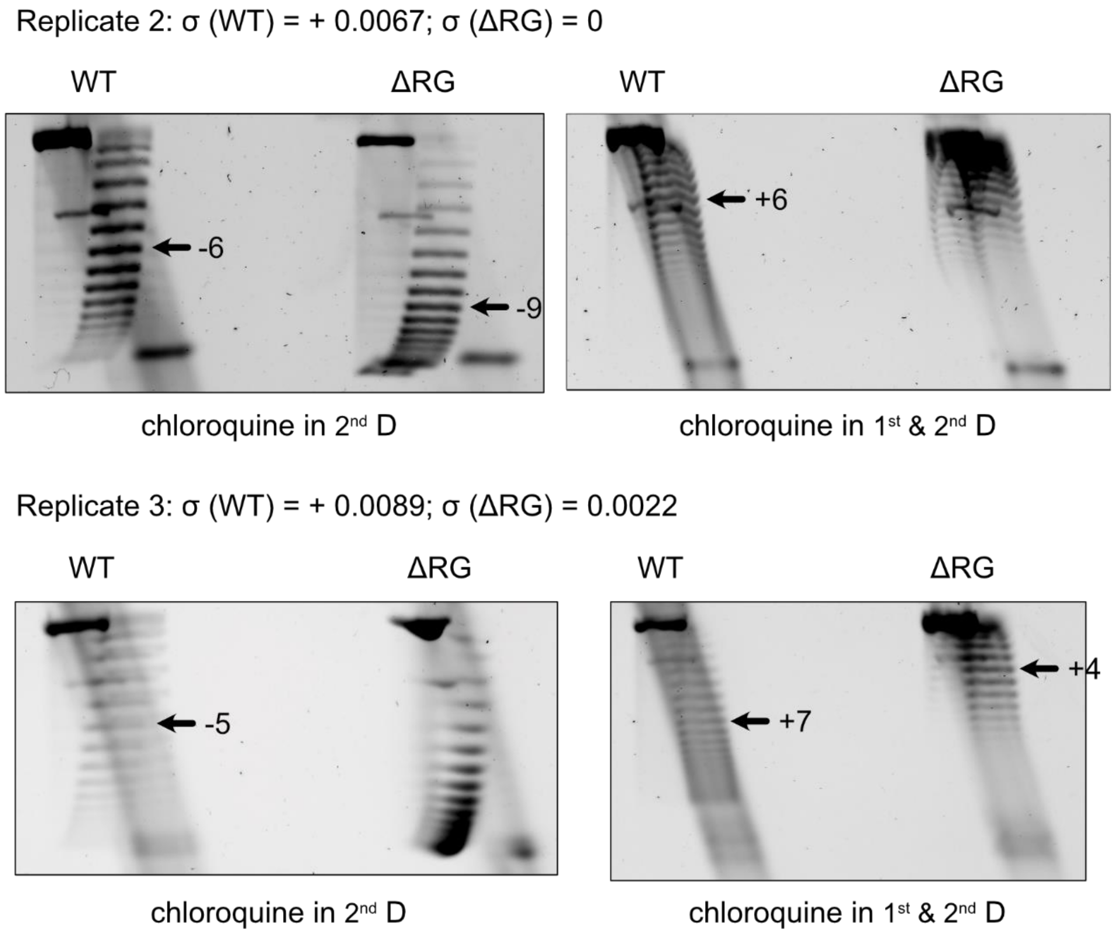


**Figure S7. Plasmid topology analysis.**

Topology of reporter plasmid pTPTK3 isolated from WT and ΔRG strains of *T. kodakarensis*. Plasmid DNA was isolated from WT and ΔRG strains and separated by 2D gel electrophoresis, which resolves plasmids according to the number and handedness of supercoils. For each analysis two gels were run in parallel, one where chloroquine was added only in the 2$^{nd}$ dimension (left panels) and a second where chloroquine was added both in 1$^{st}$ and 2$^{nd}$ dimensions (right panels). This latter condition allows to relax strongly negatively supercoiled plasmids thus facilitating the identification of the major topoisomers (indicated by an arrow). The superhelical density (σ) of the major topoisomer was calculated by considering, if necessary, the contribution of chloroquine that introduces positive supercoils and temperature shift (from 85°C to ~22°C) which introduces negative supercoils (see material and methods for details).

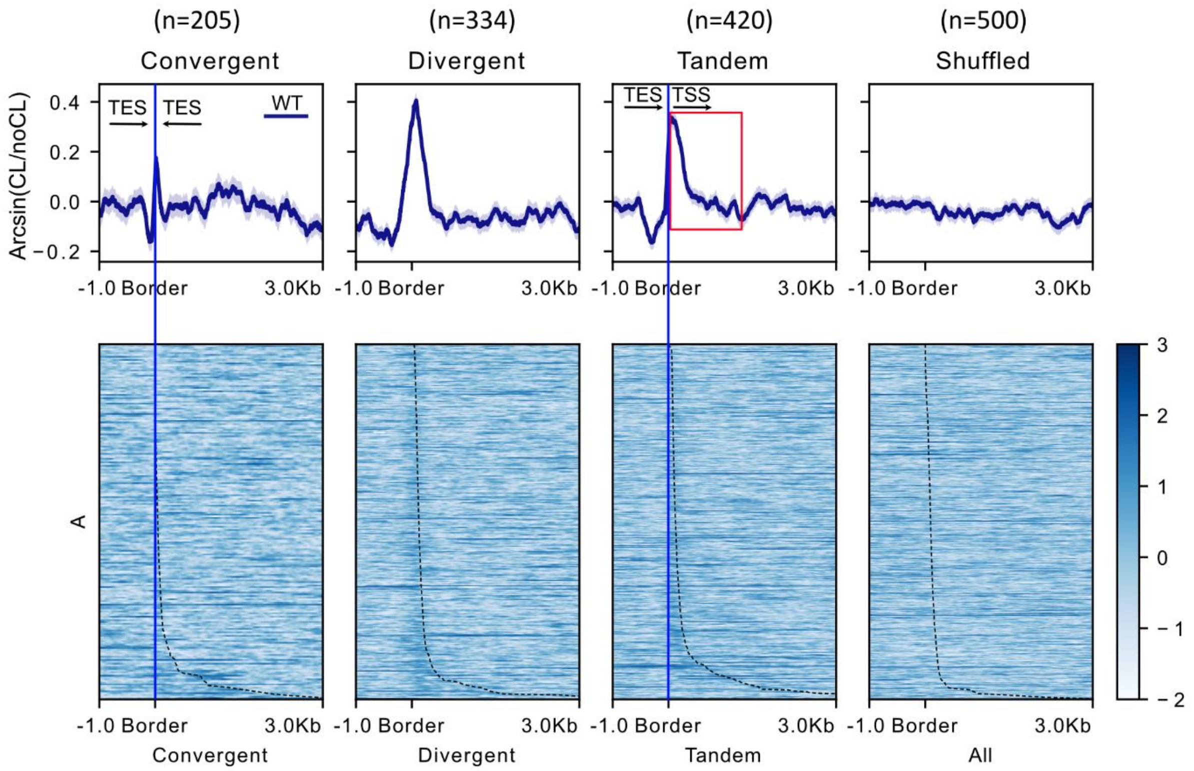


**Figure S8. Overlap in CLL signals of adjacent TSS and TES regions.**

Distribution of CLL across convergent, divergent and tandem TUs. The plots' reference point is the border of the intergenic region while the dotted line going through the heatmap represents the intergenic region length. Shuffled regions (n=500) with a length distribution matching that of intergenic regions and randomly placed across the genome were used as a control (rightmost panels). The heat maps show relative levels of CLL for each TU in a given group (one per row). The bar on the right of the heat maps color-codes the signal intensity. The two cases of intergenic regions containing TES (convergent and tandem TUs) are highlighted. Most of the signal downstream of TES (red rectangle) comes from the nearby TSS.

**Table S1. Strains and plasmids used in this study.**

| Strain name | Genotype | Markers | Plasmid | Source |
|---|---|---|---|---|
| *Escherichia coli* XL1-Blue | *endA*1 *gyrA*96 thi-1 *recA*1 *relA*1 lac *glnV*44 F'[ ::Tn10 *proAB+ lacIq* Δ(l*acZ*)M15] *hsdR*17 | Tetracycline resistance Nalidixic acid resistance | / | Stratagene |
| *T. kodakarensis* TS559 | Δ*pyrF*; Δ*trpE::pyrF*, Δ*TK0664*, Δ*TK0149* | Uracil prototrophy Tryptophan auxotrophy Agmatine auxotrophy 6-methylpurine resistant | / | Santangelo et al., 2010 |
| *T. kodakarensis* DAD | Δ*pyrF, ΔTK0149, ΔTK0143** | Uracil prototrophy Agmatine auxotrophy | / | Hiroki Higashibata |
| *T. kodakarensis* DADΔRG | Δ*pyrF, ΔTK0149, ΔTK0143*, ΔTK0470* | Uracil prototrophy Agmatine auxotrophy | / | Hiroki Higashibata |
| *T. kodakarensis* TKDAD | Δ*pyrF*; Δ*trpE*::*pyrF*, Δ*TK0149, ΔTK0143* | Tryptophan auxotrophy Agmatine auxotrophy | / | This study |
| *T. kodakarensis* TKΔRG | Δ*pyrF*; Δ*trpE*::*pyrF*, Δ*TK0149, ΔTK0143* | Tryptophan auxotrophy Agmatine auxotrophy | / | This study |
| *T. kodakarensis* TKDAD*p* | Δ*pyrF*; Δ*trpE*::*pyrF*, Δ*TK0149, ΔTK0143* | Agmatine auxotrophy | pTPTK3 | This study |
| *T. kodakarensis* TKΔRG*p* | Δ*pyrF*; Δ*trpE*::*pyrF*, Δ*TK0149, ΔTK0143* | Agmatine auxotrophy | pTPTK3 | This study |
| *T.kodakarensis* TKAg | Δ*pyrF*; Δ*trpE::pyrF*, Δ*TK0664*, Δ*TK0149* | Uracil prototrophy Tryptophan auxotrophy 6-methylpurine resistant | pTNAg | Villain et al., 2021 |
| *T. kodakarensis* TKY119F | Δ*pyrF*; Δ*trpE::pyrF*, Δ*TK0664*, Δ*TK0149* | Uracil prototrophy Tryptophan auxotrophy 6-methylpurine resistant | pTNAg-Y119F | Villain et al., 2021 |
| *T. kodakarensis* TKgyrAB | Δ*pyrF*; Δ*trpE::pyrF*, Δ*TK0664*, Δ*TK0149* | Uracil prototrophy Tryptophan auxotrophy 6-methylpurine resistant | pTNAg-gyrAB | Villain et al., 2021 |
| **Plasmid name** | **Genotype** | ***E. coli* marker(s)** | ***Thermococcus* marker(s)** | **Source (Accession No.)** |
| pTPTK3 | pTP2::(p15A-*cat*),(PTK0149-*TK0149*) | CmR | Agm | Catchpole et al., 2018 (MG920816) |

| | | | | |
|---|---|---|---|---|
| pTNAg | pLC70Δ(*TK0254-PF1848*):: (PTK0149-*TK0149*) | AmpR, KanR | Agm | Catchpole et al., 2018 (MG920813) |
| pLC70 | see reference | AmpR, KanR | Trp, MevR | Santangelo et al., 2008 (N/A) |
| pTNAg-gyrAB | pLC70Δ(*TK0254-PF1848*):: (PTK0149-*TK0149*), PhmtB-gyrAB | AmpR, KanR | Agm | Villain et al., 2021 |
| pTNAg-Y119F | pLC70Δ(TK0254-PF1848):: (PTK0149-TK0149), PhmtB-gyrAB(Y119F) | AmpR, KanR | Agm | Villain et al., 2021 |

*TK0134 (TK_RS00660) is annotated as probable acetylpolyamine aminohydrolase, histone deacetylase family (GenBank);

**Table S2. Differential expression of genes with functions in DNA repair and DNA topology**

| | | **TKΔRG vs TKDAD** | | | |
|---|---|---|---|---|---|
| Locus (old/new) nomenclature | Annotation | log2FC | $P_{adj}$ | NC (TKDAD) | NC (TKΔRG) |
| **DNA topoisomerases** | | | | | |
| *TK0778/ TK_RS03845* | **Mini A*** | **0.38** | **3.01e-06** | 993.5 | 1592.5 |
| *TK0798/ TK_RS03950* | DNA topoisomerase VI, subunit A | -0.02 | 0.87 | 1438.5 | 1737.5 |
| *TK0799/ TK_RS03955* | DNA topoisomerase VI, subunit B | -0.05 | 0.59 | 1685.3 | 1983.25 |
| *TK1091/ TK_RS05360* | **DNA topoisomerase III** | **0.37** | **8.02e-06** | 1064.8 | 1671.5 |
| **NAPs and histones** | | | | | |
| *TK1413/ TK_RS07015* | Histone A | -0.16 | 0.28 | 2115.5 | 2311,5 |
| *TK2289/ TK_RS11530* | Histone B | 0.22 | 0.03 | 1360 | 1952.8 |
| *TK0471/ TK_RS02325* | **TrmBL2** | **-0.64** | **3.4e-07** | 3151.3 | 2462.5 |
| *TK0560/ TK_RS02760* | Alba | 0.05 | 0.57 | 5088.8 | 6518.5 |
| *TK1040/ TK_RS05120* | Histone-fold protein | **-0,80** | **2.25e-07** | 2372.5 | 1639 |
| *TK0750/ TK_RS05120* | Histone-fold protein | 0.11 | 0.46 | 449.3 | 600 |
| **DNA repair - MMR** | | | | | |
| *TK0682/ TK_RS03370* | MutS-like | -0.94 | 0.42 | 540.8 | 611.5 |

| | | | | | |
|---|---|---|---|---|---|
| *TK1898/ TK_RS09500* | NucS | 0.08 | 0.42 | 345 | 446 |
| *TK1175/ TK_RS05785* | Hjc | 0.004 | 0.98 | 116.5 | 142.5 |
| **DNA repair - HR** | | | | | |
| *TK1332/ TK_RS12245* | **Hel308** | **0.56** | **1.5e-10** | 903.3 | 1621.3 |
| *TK2212/ TK_RS11115* | **Mre11** | **-0.94** | **2.9e-18** | 577.3 | 362.5 |
| *TK2210/ TK_RS11105* | **NurA** | **-0.77** | **1,4e-07** | 481.3 | 339.8 |
| *TK2211/ TK_RS11110* | **Rad50** | **-1.69** | **5.1e-44** | 940 | 351.3 |
| *TK1899/ TK_RS09505* | RadA | 0.29 | 2.9e-05 | 5775.5 | 8734.3 |
| **DNA repair - NER** | | | | | |
| *TK1281/ TK_RS06335* | Fen1/Xpg | 0.02 | 0.83 | 911.25 | 1129.5 |
| *TK1021/ TK_RS05025* | **Hef/Xpf** | **0.36** | **0.00022** | 1007.75 | 1580.25 |
| *TK0928/ TK_RS04570* | **Xpb** | **-0.66** | **2,91e-15** | 1148.25 | 879.5 |
| *TK0784/ TK_RS03875* | Xpd | -0.13 | 0.21 | 640.25 | 710.25 |
| **DNA repair - BER** | | | | | |
| *TK0345/ TK_RS01700* | AlkA | 0.15 | 0.18 | 402.25 | 545.25 |
| *TK0940/ TK_RS04625* | **8-oxoguanine DNA glycosylase** | **1.52** | **3.25e-36** | 227.5 | 798.25 |
| *TK2140/* | Lig1 | -0.06 | 0.46 | 1348.75 | 1574.75 |

| | | | | | |
|---|---|---|---|---|---|
| *TK_RS10745* | | | | | |
| *TK0170/ TK_RS00835* | Nfo/Endonuclease IV | -0.23 | 0.27 | 827.25 | 855.5 |
| *TK1141/ TK_RS05625* | NTH/Endonuclease III | -0.12 | 0.5 | 348.5 | 392.25 |
| *TK1252/ TK_RS06185* | RecJ | -0.08 | 0.41 | 1053.25 | 1201 |
| **Replication protein A (RPA)** | | | | | |
| *TK1959/ TK_RS09810* | **RPA31** | **1.69** | **4,27e-67** | 990.5 | 3918 |
| *TK1960/ TK_RS09815* | **RPA14** | **1.91** | **1,57e-79** | 517.25 | 2373,75 |
| *TK1961/ TK_RS09820* | **RPA41** | **1.93** | **2,33e-69** | 808.25 | 3785.25 |

NC (TKDAD) – normalised counts TKDAD strain; NC (TKΔRG) normalised counts TKΔRG strain; mean value of four replicates is given. The values above the significance threshold (Padj ≤ 0.01 and $Log_2FC \geq |0.33|$) are indicated in bold letters. UniProt and original publications (Gehring et al., 2020; Fujikane et al., 2010) were used as sources for annotation of known DNA repair genes.

* homolog of the A-subunit of Topo VIII (Takahashi et al., 2020)
